# Large language model-based bibliometric evaluation of population descriptors in human genetics

**DOI:** 10.64898/2026.09.24.754051

**Authors:** Stephanie M. Gogarten, Daphne O. Martschenko, Roshni Patel, Nasa Sinnott-Armstrong

## Abstract

As the use of population descriptors such as race, ethnicity, and ancestry have become increasingly common in modern genetics research, there have been growing calls to critically examine their use. Most notably, in 2023, the National Academies of Science, Engineering, and Medicine (NASEM) published a report titled *Using Population Descriptors in Genetics and Genomics Research: A New Framework for an Evolving Field*, which included eight specific and actionable recommendations for researchers to implement the ethical and accurate use of population descriptors in genetic research.

Here, we use the 2023 NASEM report as a benchmark to analyze the use of population descriptors in genome-wide association studies (GWAS). We develop a general toolkit for large language model-based bibliometrics, operationalize the report’s recommendations into an evaluation framework, and apply this framework to evaluate all 4,007 papers from the GWAS Catalog published between 2007 and 2025 with full text available on PubMedCentral. We find significant improvements in adherence to NASEM report recommendations over time. However, most improvements predate the publication of the NASEM report itself, suggesting the report functioned primarily as a synthesis of existing best practices rather than a catalyst for change. We conclude by highlighting opportunities for growth in the field of human genetics.

## Introduction

The use of population descriptors such as race, ethnicity, and genetic ancestry in biomedical research has grown substantially over the past two decades. Biomedical researchers have long used population descriptors to characterize research participant diversity and study health disparities. The genomic era has additionally led to broader use of population descriptors to identify and adjust for population structure in genome-wide association studies (GWAS). Though population descriptors can provide some utility in controlling for population structure and other types of confounding, there are several concerns associated with their use (Callier 2019; Brothers et al. 2021; Khan et al. 2022). In particular, conflation of race and ethnicity with genetics—and more generally, conflation of discrete categories with continuous genetic variation—risks contributing to bioessentialism. Throughout a number of genetic studies, there is evidence that these simplifying assumptions can undermine scientific rigor and reproducibility, and reinforce harmful misconceptions about human genetic diversity (CITE).

In 2023, the National Academies of Science, Engineering, and Medicine published a report titled *Using Population Descriptors in Genetics and Genomics Research: A New Framework for an Evolving Field* (*Using Population Descriptors in Genetics and Genomics Research* 2023). The report, which emphasizes the importance of accurate, ethical, transparent, and justified use of population descriptors, was the culmination of a multi-year interdisciplinary effort spanning geneticists, social scientists, historians, and members of the public. Briefly, the report recommended that researchers differentiate between race, ethnicity, and genetic ancestry, avoid typological thinking, engage with communities, use thoughtful language, and be clear and consistent in applying population labels. In the three years since publication, the NASEM report has been cited over 250 times by geneticists, journal editors, and ELSI scholars as an important reference on the roles and responsibilities associated with ethical use of population descriptors.

Here, we use the NASEM report as a benchmark to comprehensively and quantitatively analyze the use of population descriptors in GWAS publications. To do so, we pilot the use of a large language model (LLM)-based tool for bibliometrics in biomedical research, enabling bespoke, systematic bibliometric analysis on an unprecedented scale. While some studies have developed automated tools for text mining biomedical research (González-Márquez et al. 2024; Bifarin et al. 2025; Vos et al. 2026), most focus solely on abstracts, and none have developed tools designed specifically for GWAS publications. Leveraging our genetics-focused toolkit, we characterize which population descriptors have been used in GWAS and evaluate how well they adhere to the NASEM report recommendations. We conclude by synthesizing the field’s history of population descriptor reform and discussing opportunities for the future use of LLMs in bibliometrics of biomedical research.

## Methods

### Preprocessing of GWAS papers

We first identified papers from the GWAS catalog published between March 5, 2007 and December 5, 2025 for which full text in XML format was available from PubMedCentral (Welter et al. 2014; MacArthur et al. 2017; Sollis et al. 2022). In total, this yielded 4,007 papers. For each paper, we extracted the section headers and body text, omitting the author list, tables, figures, and references.

To determine the sample size for papers, we used the maximum reported sample size for papers that included multiple individual GWAS datasets. To assess the ancestral category and country of recruitment for each paper, we downloaded the study metadata from the GWAS Catalog, which included these fields. Most papers had multiple studies listed, each with a Broad Ancestral Category following the HANCESTRO ontology (Morales et al. 2018) and a Country of Recruitment. To analyze trends in ancestral category and country of recruitment, we filtered all studies to only include discovery cohorts and not replication samples.

### Developing a framework for large language model-based bibliometrics

To evaluate the use of population descriptors in GWAS papers, we created a LLM prompt that included the text of the eight recommendations of the NASEM report (*Using Population Descriptors in Genetics and Genomics Research* 2023) as evaluation criteria. We identified example text from two papers, one of which we judged as following the recommendations very well (Belbin et al. 2021), and another paper that we judged as demonstrating typological thinking and failing to follow the recommendations (Karol et al. 2017). We included this example text in the prompt along with instructions that the former should be given a rating of 5, while the latter should be given a rating of 1 (see Supplementary Material for the full text of the prompt). This series of structured questions and examples followed best practices for LLM prompt design, and enabled us to reproducibly quantify relevant language in the text of each manuscript (Colangelo et al. 2025).

To interact with the LLMs, we used the ellmer R package (Wickham et al., n.d.). After providing the initial prompt, we asked the LLM the following question: “Please provide an overall rating of how well the paper follows all eight of the recommendations on a scale of 1 to 5. If the paper does not mention any population descriptors, respond with rating ‘0’.” If the agent responded with a rating > 0, we then asked for a rating corresponding to each of the eight recommendations (i.e. “Please evaluate how well the paper follows recommendation X on a scale of 1 to 5”). Finally, we asked: “Please list all population labels mentioned in the paper.” We instructed the LLM to respond with results in JSON format, which we then parsed in R. Malformed entries (n = 3 out of 4,007 across runs) were excluded.

To analyze the population descriptors contained in each GWAS paper, we parsed the LLM-outputted population labels to identify descriptors containing the capitalization-invariant substring “ancestry,” as well as other terms including demonyms and racial and ethnic categories. We additionally conducted a simple lowercase text match search to identify papers containing the term “trans-ethnic”; this use of string searching was necessary because “trans-ethnic” is not itself a population descriptor.

### Evaluating large language models

To evaluate the performance of different LLMs, we first selected a random subset of 10 papers. One of us (SMG) read each of the 10 papers, assigning ratings for how well the paper followed each of the recommendations separately, how well it followed the recommendations overall, and which population descriptors were used in the paper.

Using the prompt framework described above, we compared four LLMs: Llama3, Google gemini-2.5-flash, Google gemini-2.5-pro, and Perplexity sonar-reasoning-pro. Llama3 responses were inconsistent with human ratings and prone to hallucinations, so it was excluded from further analyses. While Perplexity was broadly consistent with other models (Supplementary Figure 1), it did not always return results in a parseable format, and moreover had the tendency to become stuck in a seemingly infinite reasoning loop where it reconsidered its answer over and over again.

**Figure 1.**
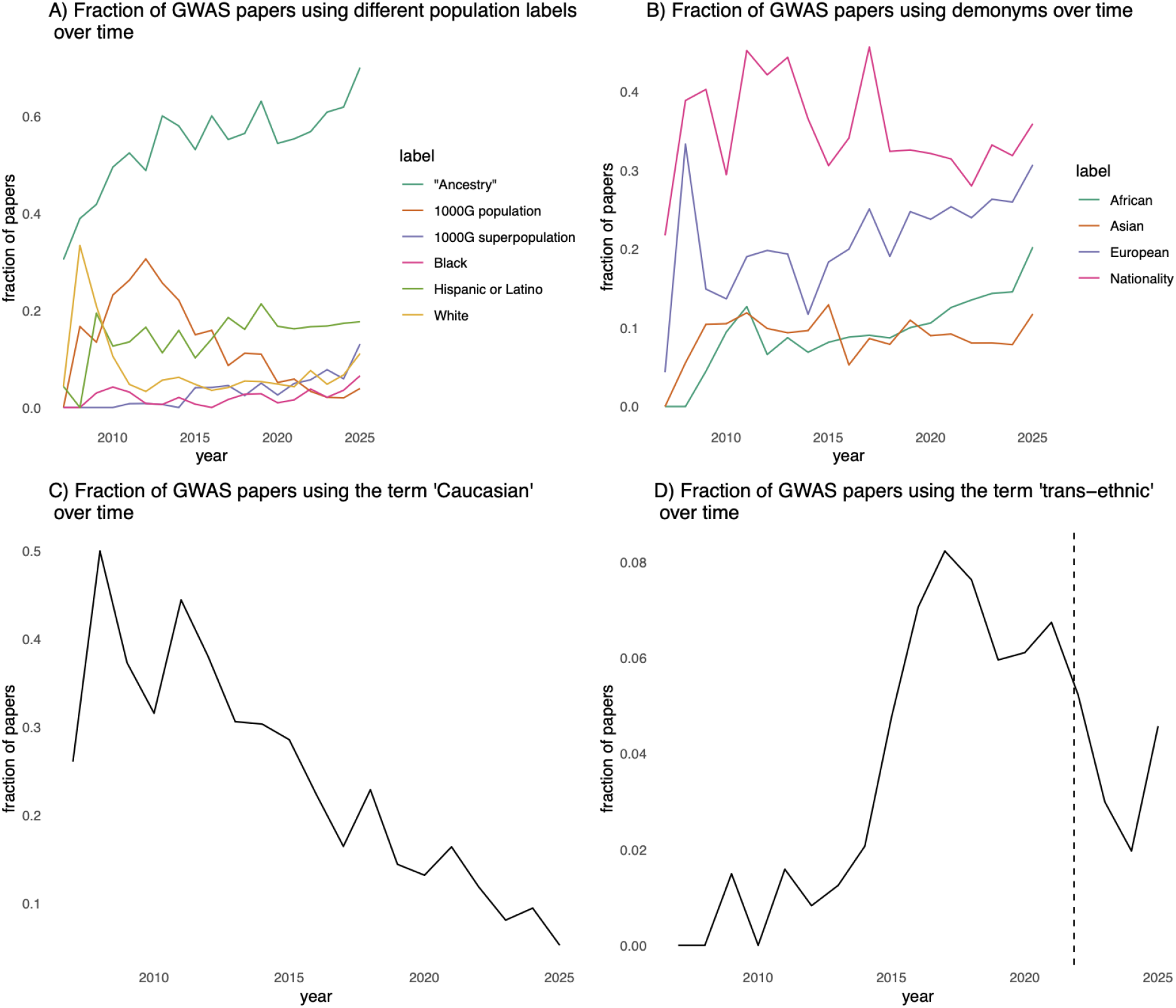
A) Fraction of GWAS papers using the population labels “-Ancestry”, any 1000G population (e.g. CEU, YRI), and 1000G superpopulation (e.g. AFR, EUR), and the racial/ethnic labels “Black”, “White”, “Hispanic” and “Latino” by year. B) Fraction of GWAS papers using demonyms (e.g. “British”, “American”, “African”) as population labels by year. C) Fraction of GWAS papers containing the term “Caucasian” by year. D) Fraction of GWAS papers containing the term “trans-ethnic” by year. The vertical line is the publication date of Kamariza et al, “Misuse of the term ‘trans-ethnic’ in genomics research.”

We thus proceeded with comparing the two Google models for all papers, asking each model to return an overall rating for each paper. To assess reproducibility, we ran gemini-2.5-flash and gemini-2.5-pro on the same sample of papers twice each (Supplementary Figure 2). We found results were fairly consistent (mean correlation = 0.68), especially when comparing “high” ratings (4 or 5) to “low” ratings (1 or 2) (Supplementary Figure 3). Given the consistency of results and that gemini-2.5-flash was significantly faster, we chose this model to use for subsequent queries.

**Figure 2.**
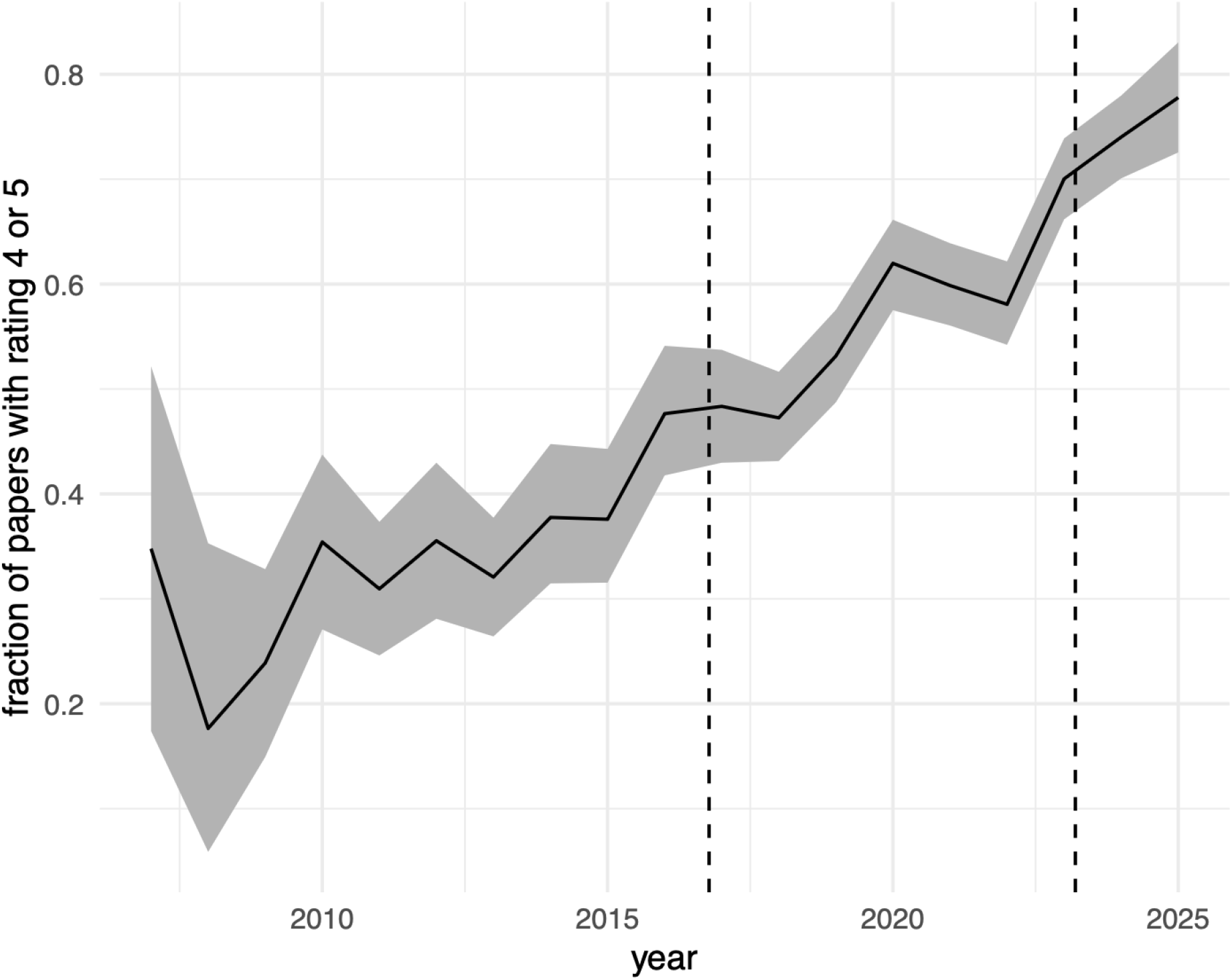
Fraction of high ratings (4 or 5) by year. Error bars are estimated from bootstrapping 1000 replicates. Vertical dashed lines are publication dates of Popejoy and Fullerton, “Genomics is failing on diversity” (2016) and the NASEM report (2023).

**Figure 3.**
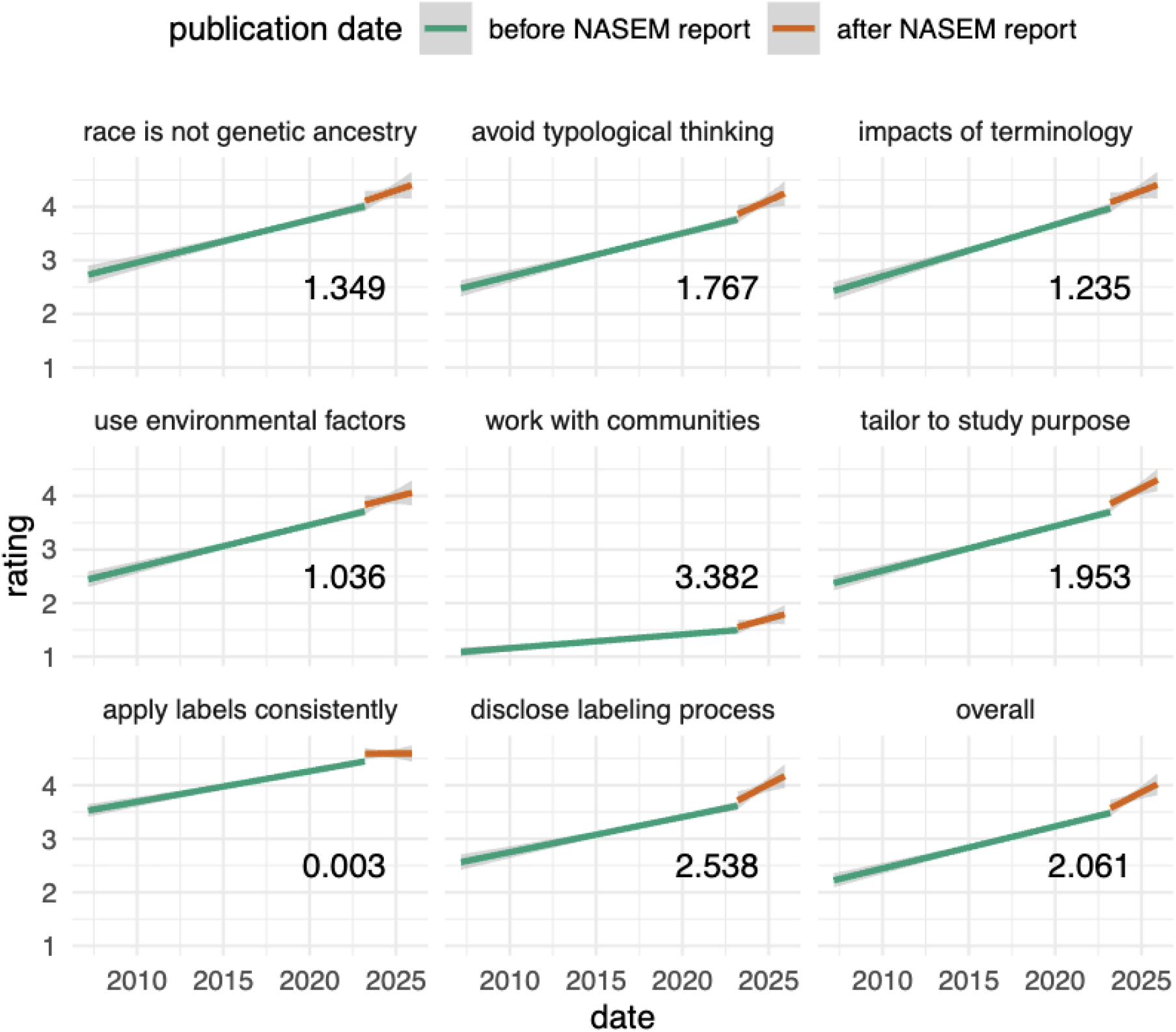
Linear model fits of rating over time, before and after the publication of the NASEM report. The number displayed in each panel is the ratio of the slopes after the NASEM report versus before, and panels correspond to each of the NASEM recommendations (and/or the overall score across recommendations).

We found that gemini-2.5-flash was broadly consistent with human-assigned ratings (mean correlation = 0.58), especially given the difficulty and perceived subjectivity in assigning human ratings for many of the papers (Supplementary Figure 4). The most significant difference was that for two papers, the human assessment was that the papers did not use descent-associated population descriptors, while the LLM proceeded to assign ratings to both papers. In one of these cases (Gu et al. 2011), the paper received a low rating of 2 with the reasoning that it failed to report on population descriptors (Supplementary Results, example 1). In the other case(Nounu et al. 2021), the paper received a high rating of 4 since it used principal components to account for genetic ancestry and avoided problematic terminology (Supplementary Results, example 10).

**Figure 4.**
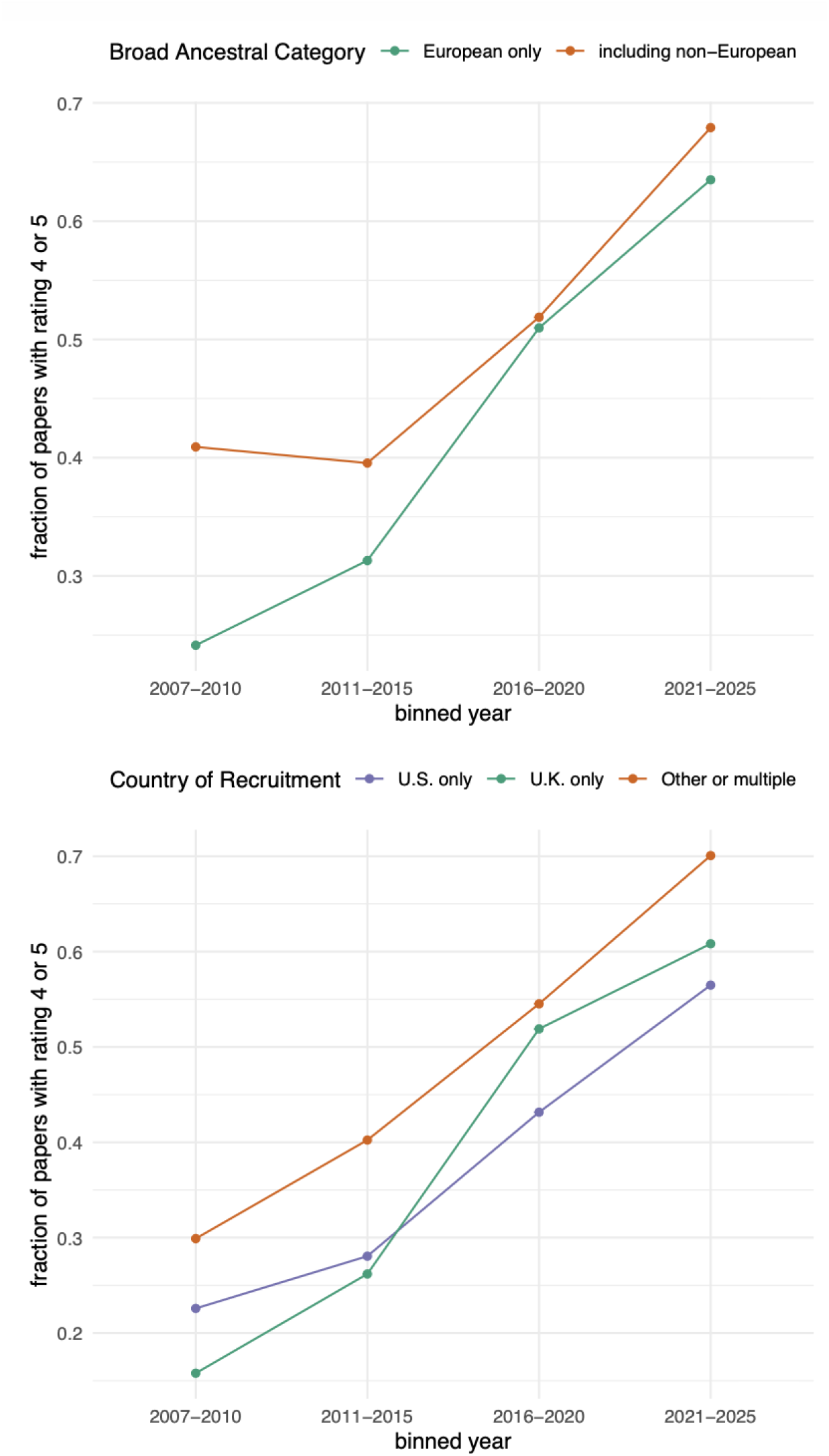
A) Ratio of high ratings (4 or 5) by year, split by Broad ancestral category of study participants as entered in the GWAS Catalog. Categories are “European” for studies with only European ancestry participants (n = 2059), and “Other or multiple” for all other studies (n = 1696). B) Fraction of high ratings (4 or 5) by year, split by country of recruitment. Categories are “U.S” where all participants are from the U.S. (n = 468), “U.K.” where all participants are from the U.K. (n = 561), and “Other or multiple” for all other studies (n = 2256).

As an additional evaluation metric, we compared LLM-based identification of the population descriptor “Caucasian” against a simple exact string match in the full text of each paper (Supplementary Figure X). The LLM accurately identified the term in 726 papers and correctly determined its absence in 3,268 papers. In total, the LLM had 12 false negatives and one false positive. We additionally evaluated whether papers containing the term “Caucasian” received a low rating (1-2) for recommendation 3, which explicitly cautions against the use of this term, stating that it was “originally coined to convey white supremacy” (*Using Population Descriptors in Genetics and Genomics Research* 2023). Consistent with our expectation, nearly all papers containing the term “Caucasian” received low ratings for recommendation 3 (Supplementary Figure 5).

## Results

Our dataset consisted of all papers from the GWAS catalog with open-access full text, totaling 4,007 papers. As expected, the number of papers in the GWAS catalog exhibits a supralinear increase over time, with the caveat that a number of papers published recently (i.e. in 2024 and 2025) do not yet have open-access text (Supplementary Figure 6). The majority of these papers referenced GWAS with fewer than 10,000 participants, though some GWAS had over 1,000,000 participants (Supplementary Figure 7). The journals associated with these GWAS varied, with the most common journals being Nature Communications (n = 437), Nature Genetics (n = 299), PLOS One (n = 292), and PLOS Genetics (n = 256) (Supplementary Figure 8). The most common country of primary affiliation for the first author was the U.S. (n = 1,461), followed by the U.K. (n = 492) and China (n = 333) (Supplementary Figure 9).

### Characterizing which population descriptors are used in GWAS

To analyze the use of population descriptors in each GWAS publication, we developed a LLM-based framework for bibliometric analysis (see Methods). We first sought to understand which descent-associated population descriptors were used in GWAS publications. To do so, we prompted the LLM to first identify whether a publication contained any population descriptors, and then to extract the exact phrase(s) associated with those descriptors. 3,913 publications, or 98% of our data, were detected to contain one or more population descriptors.

We found that the use of the term “ancestry” in population descriptors has increased substantially over time, from fewer than 40% of papers in 2007 to greater than 60% in 2025 (Figure 1A). Researchers frequently use “ancestry” as a suffix for geographic regions, e.g. “European ancestry” or “Chinese ancestry”, and accordingly, we also found that the use of geographic regions in population descriptors has increased over time. Among continental regions, “African” and “European” have both increased over time, while use of “Asian” increased in the late 2000s and has continued to be used in approximately 10% of papers since. Unsurprisingly, we found that “European” was much more common than “African” or “Asian”, reflecting the demographics of GWAS study participants (Popejoy and Fullerton 2016).

As ancestry-related terms have increased in frequency, use of race and ethnicity has exhibited a parallel decrease. Use of the racial term “White” has decreased since the late 2000s, while use of the term “Black” remains low but has shown a modest increase in the past five years. Use of the terms “Hispanic” and “Latino” has increased since the late 2000s and has remained steady over the past decade.

Surprisingly, our results also suggest a marked shift towards broader, less specific population descriptors in GWAS publications over time. This practice conflicts with recent recommendations to use narrower, more specific descriptors (Lewis et al. 2022), and may reflect the rapid expansion and diversification of biobanks. Use of 1000 Genomes population labels (e.g. “CEU”, “YRI”) peaked in the early 2010s and has declined since. Meanwhile, use of 1000 Genomes superpopulation labels (e.g. “AFR”, “EUR”) has increased over time, and overtook the use of narrower population labels after ∼2021. Use of nationality labels (e.g. “British”, “American”) was highest in the early 2010s, and has declined since.

Finally, we analyzed the frequency of problematic population descriptors that have been previously criticized by high-profile publications. Specifically, we analyzed the use of “Caucasian”, which was criticized by Moses (2017) and Popejoy (2021); and the use of “trans-ethnic”, which was criticized by Kamariza et al. (2021). The use of both terms has declined in recent years: “Caucasian” declined from over 50% of GWAS publications in 2008 to less than 10% in 2023, while use of “trans-ethnic” initially increased, peaked in 2017, and has subsequently declined (Figure 3C, 3D). Notably, in both cases, the decline of the term was well underway by the time of publications criticizing its use, potentially reflecting shifting field norms that preceded the publications in question.

### Evaluating how population descriptors are applied in GWAS

After assessing which population descriptors were used in GWAS publications, we next sought to evaluate *how* these descriptors were applied. Specifically, we developed an evaluation rubric centered on the eight recommendations on the use of population descriptors published in the 2023 NASEM report. Using this rubric, we prompted the LLM to rate each GWAS publication on how well it followed each individual recommendation, on a scale of 1 (very poor) to 5 (very well) (see Supplement for the full prompt text). We additionally prompted the LLM to give each publication an overall rating, ranging from 1 to 5.

To analyze how well GWAS publications adhered to the NASEM recommendations, we grouped publications into high-scoring (ratings of 4 or 5) and low-scoring (ratings of 1 or 2) categories for each recommendation and for the overall rating. We used these rating categories because while numeric ratings varied across LLM replicates, these broader rating categories remained consistent (Supplementary Figure 3). In total, 2,088 papers were categorized as high-scoring, 1,322 papers were categorized as low-scoring, and 503 papers received a rating of 3.

To understand broad trends in the use of population descriptors, we first evaluated the overall rating assigned to every GWAS publication (Figure 1, Supplementary Figure 10). We observed a clear improvement in overall adherence to the NASEM recommendations over time, as evidenced by an increasing proportion of high-scoring papers (beta = 0.088 per year, p < 2e-16). Moreover, the rate of improvement appears to have accelerated in the past decade (0.095 per year, p < 2e-16) relative to the years 2007 - 2015 (0.045 year, p = 0.035). Perhaps not coincidentally, the field demonstrated a growing awareness of the importance of diversity in GWAS around this time, notably with the publication of Popejoy and Fullerton’s “Genomics is failing on diversity” in 2016 (Popejoy and Fullerton 2016).

We next characterized adherence to individual recommendations (Figure 2, Supplementary Figures 11-18). We found that GWAS publications received varying ratings across individual recommendations (Supplementary Figure 19). Overall publication ratings were highly correlated with recommendations 1 (“race is not genetic ancestry”), 2 (“avoid typological thinking”), 3 (“impacts of terminology”), and 6 (“tailor to study purpose”). In contrast, recommendation 5 (“work with communities”) was the least correlated with the overall rating and other recommendations, and also exhibited the lowest ratings overall (mean = 1.41).

Finally, we analyzed the impact of the NASEM report on how population descriptors are applied. To do so, we fit separate regression models for the time period before and after the publication of the NASEM report (March 14, 2023). For most recommendations, we found that the ratio of the post-NASEM slope to the pre-NASEM slope is greater than 1, indicating that uptake of the recommendations increased after the publication of the NASEM report. One exception was the recommendation to apply labels consistently, which had the highest rating of any of the recommendations and a nearly flat slope after the publication of the NASEM report. Interestingly, though the recommendation to engage with communities had the lowest ratings overall, it exhibited the largest increase in slope after the NASEM report was published (pre-NASEM slope = 0.025 per year; post-NASEM slope = 0.061 per year).

### Analyzing the role of publication demographics

Given that the NASEM report articulates how to apply population descriptors to datasets, we hypothesized that adherence to NASEM recommendations might vary with dataset demographics. We first evaluated the role of the ancestral category associated with each GWAS. Because GWAS are predominantly biased toward European ancestries (Sirugo et al. 2019), we divided our dataset into two categories: papers which only included GWAS annotated as “European” (n = 2,059), and all other papers (n = 1,696). To improve power to detect a difference between the two categories, we binned publications into four time periods of approximately 5 years each.

We found that papers including GWAS on non-European individuals consistently demonstrated better adherence to NASEM recommendations, especially in years 2007-2015 (Figure 4A). In more recent years, the proportion of high-scoring papers among GWAS including these populations has remained higher, but the gap has narrowed.

We next evaluated the role of the country of dataset recruitment. We found that the most common countries were the United States (n = 468) and the United Kingdom (n = 561). Echoing our analysis of ancestral category, we found that GWAS publications with datasets exclusively from the U.S. and U.K. had a smaller proportion of high-scoring papers relative to publications with datasets from other countries (Figure 4B). We additionally evaluated the role of the country of the first author on the publication for countries with > 100 papers (Supplementary Figure 20). We found that adherence to NASEM recommendations varied by country of affiliation, with no significant trends.

## Discussion

The fields of genetics and genomics have expanded dramatically in the past twenty years, including larger datasets, more publications, and broader applications. Bibliometric analyses offer an opportunity to evaluate practical, ethical, and methodological questions across this growing literature. Here, we conduct what is, to our knowledge, the first use of LLMs for bibliometric analysis of the human genetics literature. Using this toolkit, we characterize and evaluate the use of population descriptors across the GWAS Catalog, demonstrating the utility of LLMs for large-scale bibliometric investigation. Moreover, our work underscores the possibility of bibliometrics as a tool for reflexivity: quantifying population descriptors and other methodological decisions can help identify best practices and areas for growth, especially in a rapidly developing field.

We found that the choice of LLM was crucial to the success of the project. Among the models we tested, their ability to reliably return accurate and parseable results varied tremendously. Building infrastructure to support more reliable reasoning and mitigate failures such as indecision paralysis will be important for scaling the use of LLMs for scientific tasks. Another difficulty with use of LLMs accessed by API is that they are subject to change by the provider at any time. One model showed initial promise in matching human ratings, but the provider updated the model and subsequently it no longer performed well. Building open model standards and persistent access to versioned LLMs is a critical component of building reproducible results on these platforms.

We also note that LLMs are not deterministic. As described in the Methods, they do not produce exactly the same response to the same question when asked multiple times on distinct instantiations. Therefore, it is important to assess consistency by comparing the results of multiple rounds of asking the same question to both the same and separate LLMs (see Supplementary Figure 2). Application of ensemble learning methods increase robustness of results, provided the additional costs and time needed to generate these results are acceptable for the given application. For any model, it is also essential to compare the results with human reviews for a subset of data in order to assess whether it is performing well. Developing strategies to incorporate human feedback into the model review will further increase accuracy of these approaches (Templin et al. 2026).

Despite these limitations, some LLMs performed remarkably well. Within seconds, the LLMs we employed were able to generate a comprehensive summary of population-descriptor use that closely matched human assessment, enabling us to both broadly characterize and critically evaluate the use of population descriptors in GWAS publications. We hypothesize that new modeling advances will continue to improve accuracy, consistency, and speed beyond the approaches available today, but additional safeguards and cautious evaluation of these tools will continue to be necessary.

Our results illustrate a complex history of population descriptors in human genetics. Longitudinal trends in the population descriptors used in GWAS reflect both increasing diversity in datasets (Popejoy and Fullerton 2016; Mills and Rahal 2019), as well as a field-wide push towards replacing racialized descriptors with measures of ancestry (Panofsky and Bliss 2017; Duello et al. 2021; *Using Population Descriptors in Genetics and Genomics Research* 2023). In evaluating how GWAS papers apply population descriptors, we identified two key trends. First, while adherence to the NASEM recommendations increases after the publication of the NASEM report, we also found that significant uptake of the recommendations precedes the NASEM report by at least a decade. These results parallel our analyses of “Caucasian” and “trans-ethnic” in Figure 1, and reiterate that the publication of the NASEM report might not itself be an inflection point.

Instead, we suggest that the NASEM report codified a set of expectations reflecting ongoing discussion within human genetics. In recent years, there have been numerous calls to reckon with the use of race and ancestry in biomedical research—and particularly in genetics—in recognition of the fact that it is both scientifically inaccurate and ethically problematic to categorize human diversity into discrete groups, particularly those defined by race and/or ethnicity. Our results demonstrate that alongside this discourse, there have been quantifiable improvements in the use of population descriptors: relative to 10 years ago, GWAS publications today are much more likely to avoid typological thinking, apply labels consistently and transparently, use thoughtful language in referencing population descriptors, and move away from conflating race with genetic ancestry.

Yet, our results also identify critical areas for growth, underscoring that ongoing work is needed to ensure meaningful change. We found that GWAS using data from non-European populations, and data from outside the U.S. and U.K., exhibit stronger adherence to NASEM recommendations. This finding is notable, particularly given that the NASEM report was commissioned and published by U.S. institutions. This pattern may reflect differences in norms and practices across datasets, research communities, and study contexts: one possible explanation is that U.S.-based datasets have been slower to adopt best practices, while another is that non-U.S. countries may have received better ratings because the U.S.-centric NASEM recommendations did not target the use of population descriptors in many global contexts. Ultimately, ensuring the accurate and ethical use of population descriptors worldwide will require recommendations that account for the varied historical, cultural, and social contexts in which these descriptors are used.

Moreover, we found that GWAS publications consistently received the lowest ratings for the recommendation to “work with communities”, with an average rating of less than 2 even in recent years. This likely reflects the lack of infrastructure within the field to meet this recommendation: while many of the NASEM recommendations pertain to choices made by researchers during data analysis and manuscript preparation, working with communities requires sustained investment beginning in the initial study design. Given the profound social impacts of human genetics and genomics research, we echo recent calls (Claw et al. 2018; Hudson et al. 2020; Fox 2020; Fatumo et al. 2022; Copeland-Hardin et al. 2026) that increasing community engagement remains of paramount concern.

Ultimately, our results highlight the importance of the ongoing discussion around ethical best practices in human genetics. The changes we observed are unlikely to be attributable to any single intervention, including the NASEM report; rather, they appear to reflect the cumulative effects of incremental shifts in research practices, community expectations, and disciplinary norms. Engaging with the ethical and social implications of human genetics at all stages of study design, implementation, and dissemination will be critical to continuing this progress and building more consistent and responsible practices for the next generation of research.

## Acknowledgements

We would like to thank Jedidiah Carlson, Tami Gjorgieva, Xiyal Mukund, and Rachel Ungar for helpful feedback and suggestions on earlier versions of this work. This project was supported by the Pew Biomedical Scholars Program, NCI R01CA273198, and NHGRI RM1HG010461 (to NS-A). We thank the participants in the numerous GWAS studies that contributed to the analyses in this work.

