## Supplement for "Large language model-based bibliometric evaluation of population descriptors in human genetics"

### GWAS ethics manuscript supplement

#### Supplementary Methods

The initial prompt provided to the chat agent was as follows:

> You are a statistical geneticist with a strong interest in Ethical, Legal and Social Implications (ELSI) of human genetics research. You have recently read the report from the National Academics of Science, Engineering and Medicine on Using Population Descriptors in Genetics and Genomics Research. This report was published recently, and was developed as a result of many papers in the field of genetics perpetuating eugenical and racist ideologies.
>
> A population descriptor is a way to classify individuals according to perceived differences among groups of people. This report focuses on descent-associated descriptors that apply when people are thought to share characteristics derived from a common origin. For example, race, ethnicity, and ancestry are descent-associated population descriptors. In contrast, a group label is a specific name used to describe a population. Examples of group labels are French (a nationality), Hispanic (an ethnicity), African (a continental geographic location), and East Asian (a geographic location).
>
> Other examples of population descriptors are genetic ancestry and genetic similarity. Genetic ancestry refers to the lines or paths through an individual's family tree by which they inherited DNA from specific ancestors. It can be useful when studying human evolutionary history. Genetic similarity, a measure of genetic resemblance among individuals, is preferred in many other contexts because it moves away from race and the misconception that humans can be grouped into discrete categories.
>
> Researchers should describe the population descriptors in the cohort(s) being studied. Population descriptors can include genetic ancestry, race, ethnicity, nationality, and geographic origin. They do not include age, sex, disease status, smoking status or medication use.
>
> The recommendations from this report are as follows:

1. Researchers should not use race as a proxy for human genetic variation. In particular, researchers should not assign genetic ancestry group labels to individuals or sets of individuals based on their race, whether self-identified or not.

2. When grouping people in studies of human genetic variation, researchers should avoid typological thinking, including the assumption and implication of hierarchy, homogeneity, distinct categories, or stability over time of the groups.

3. Researchers, as well as those who draw on their findings, should be attentive to the connotations and impacts of the terminology they use to label groups. • As an example, the term Caucasian should not be used because it was originally coined to convey white supremacy, and is often mistakenly interpreted today as a "scientific" term, thus erroneously conferring empirical legitimacy to the notion of a biological white race. • Another example of a term that should not be used is black race because it wrongly implies the existence of a discrete group of human beings, or race, who could be objectively identified as "black."

4. Researchers conducting human genetics studies should directly evaluate the environmental factors or exposures that are of potential relevance to their studies, rather than rely on population descriptors as proxies. If it is not possible to make these direct measurements and it is necessary to use population descriptors as proxies, researchers should explicitly identify how the descriptors are employed and explain why they are used and are relevant. Genetics and genomics researchers should collaborate with experts in the social sciences, epidemiology, environmental sciences, or other relevant disciplines to aid in these studies, whenever possible.

5. Researchers, especially those who collect new data or propose new courses of study for a data set, should work in ongoing partnerships with study participants and community experts to integrate the perspectives of the relevant communities and to inform the selection and use of population descriptors.

6. Researchers should tailor their use of population descriptors to the type and purpose of the study, in alignment with the guiding principles, and explain how and why they used those descriptors. Where appropriate for the study objectives, researchers should consider using multiple descriptors for each study participant to improve clarity.

7. For each descriptor selected, labels should be applied consistently to all participants. For example, if ethnicity is the descriptor, all participants should be assigned an ethnicity label, rather than labeling some by race, others by geography, and yet others by ethnicity or nationality. If researchers choose to use multiple descriptors, each descriptor should be applied consistently across all individuals in that study.

8. Researchers should disclose the process by which they selected and assigned group labels and the rationale for any grouping of samples. Where new labels are developed for legacy samples, researchers should provide descriptions of new labels relative to old labels.

> Please evaluate how well the text of a scientific paper follows these recommendations. Return a number from 0 to 5 where 0 means population descriptors were not mentioned in the text, 1 means the text reported on population descriptors such as race, ethnicity and ancestry in a way that contradicts the recommendations, and 5 means the text followed the recommendations perfectly. Also summarize how you drew this conclusion, using quotes from the text if applicable.
>
> Here is an example of text that would earn the rating 5, because it clearly differentiates between self-identified race/ethnicity and genetic ancestry: To explore the relationship between self-reported R/E and genetic ancestry, we estimated global genetic ancestry proportions for a subset of BioMe participants (N = 31,705) genotyped on the Global Screening Array (GSA). We first examined the correlation between self-reported R/E and self-reported region of origin, revealing a complex relationship between R/E and subcontinental region (Figure 1B). We then merged BioMe participants with a reference panel of 87 populations representing ancestry from seven continental or subcontinental regions. Using PCA, we demonstrate that BioMe participants represent a continuum of genetic diversity, even within self-reported R/E groups (Figure 1C). BioMe participants fall between African and non-African reference panels on principal component 1 (PC 1) and between European, Asian, American, and Oceanian reference panels on PC 2 (Figure 1D).
>
> Here is an example of text that would earn the rating 1, because it incorrectly labels racial and ethnic categories as ancestral groups and conflates them with genetic variation: Because racial disparities in relapse persisted in this trial, we also performed analyses within each of the three largest ancestral groups (white, black, Hispanic) to identify ancestry-specific variations associated with relapse.
>
> Using the recommendations and these examples, evaluate the following text. Return your output as JSON with fields “rating” and “summary”. Do not respond with any text other than the JSON output. Do not use any fields other than “rating” and “summary” in the JSON output.

After this initial prompt, we asked the following question:

> Please provide an overall rating of how well the paper follows all eight of the recommendations on a scale of 1 to 5. If the paper does not mention any population descriptors, respond with rating ‘0’.

If the agent responded with a rating $> 0$, we then asked for each recommendation (1-8):

> Please evaluate how well the paper follows recommendation X on a scale of 1 to 5.

Finally, we asked:

> Please list all population labels mentioned in the paper. Do not include any additional text in your response. Respond with a JSON array of strings.

#### Supplementary Results

We include examples of text returned by the gemini-2.5-flash LLM for 10 papers (the same set of papers that was used for comparison with human ratings).

1. PMCID 3076128. The paper receives a rating of 2. It mentions a population descriptor only once, stating in the Methods section that "Cases and controls were frequency-matched on age (±5 years), sex, and ethnicity." However, it completely fails to provide any further details as recommended by the NASEM report. The paper does not specify which ethnicity labels were used, how they were ascertained, or why ethnicity was chosen as a matching factor (violating Recommendations 6 and 8). Furthermore, the statistical analysis section mentions using "HapMap3 CEU samples" (a European ancestry reference panel) for imputation without describing the ancestry or ethnicity of the study population itself. This lack of transparency and justification for the choice of reference panel is a significant omission. While the paper avoids overtly problematic language or the direct conflation of race with genetics seen in a '1' rating, its handling of population descriptors is critically underdeveloped and lacks the transparency and justification required.
2. PMCID 3485370. The paper receives a rating of 4 because it largely adheres to the recommendations. It uses specific, self-reported ethnic labels ('southern Chinese Han ethnicity' and 'Zhuang ethnicity') rather than broad, problematic racial categories, and clearly states how this information was collected. Crucially, the study does not use these labels as a proxy for genetic variation; instead, it employs a principal component analysis, a method based on genetic similarity, to control for population stratification. The authors also directly measure and adjust for environmental factors ('alcohol intake', 'cigarette smoking'), which is a key recommendation. The text could be improved by using more nuanced language to avoid treating populations as monolithic groups (e.g., comparing 'the Chinese population' and 'the Japanese population'). While it does not explicitly discuss the distinction between self-identified ethnicity and genetic ancestry with the clarity of a perfect example, its methodological approach is sound and in line with the report's principles.
3. PMCID 4079945. The paper receives a high rating for its responsible use of population descriptors. It correctly uses a genetic ancestry descriptor ('European descent') for a clear and appropriate methodological purpose: to control for population stratification in a genome-wide association study (GWAS). The authors are transparent about this choice and describe the genetic methods used to define their cohort, such as 'population genetic approaches' and 'identical-by-state analysis and Eigenstrat'. This avoids the critical error of using race as a proxy for genetic variation (Recommendation 1). Furthermore, the use

of principal components as covariates in the analysis shows an effort to model genetic variation on a continuum. The paper avoids problematic terms like 'Caucasian' (Recommendation 3) and adjusts for relevant non-genetic factors directly (Recommendation 4). The rating is not a perfect 5 because the practice of excluding individuals who were 'determined to be admixed' or of 'non-European ancestry' creates a discrete analytical category, which is somewhat contrary to the principle of avoiding typological thinking (Recommendation 2), even though it is done for valid statistical reasons.

4. PMCID 4732768. The paper receives a rating of 2. While it correctly identifies the need to stratify analyses by population to control for genetic confounding (a standard practice in GWAS), it fails on several key recommendations from the NASEM report. Most notably, it directly contradicts Recommendation 3 by using outdated and problematic terminology. The text explicitly refers to a meta-analysis on 'replication cohorts of Caucasians', a term the report specifically advises against due to its historical use to promote white supremacy and its false implication of being a scientific or biological category. Furthermore, the paper engages in typological thinking (contradicting Recommendation 2) by grouping diverse nationalities into broad, continental categories like 'athletes of European descent' and 'participants from Africa' for meta-analysis without providing genetic data (e.g., principal component analysis) to justify these groupings or acknowledge the genetic continuum. The paper makes a valid attempt to match cohorts by geographic origin, but its language and analytical groupings are not aligned with modern best practices for describing human populations in genetics research.
5. PMCID 5715013. The paper receives a rating of 2. While it commendably avoids using "race" as a descriptor, opting instead for geographically-based ancestry labels such as "European-ancestry" and "African-ancestry" (partially following Recommendations 1 and 3), it falls short on several key recommendations. Most critically, the paper violates Recommendation 8, as it fails to disclose the process by which it assigned these group labels to participants. This lack of transparency is a significant methodological omission. Additionally, the paper's rigid separation of the cohort into two distinct continental groups for analysis (e.g., "In our GWAS of 586,626 European parental lifespans..." versus the study of participants "with African ancestry") without any discussion of the continuous nature of genetic variation constitutes a form of typological thinking, which Recommendation 2 warns against. Although analyzing genetically similar groups separately can be methodologically necessary in GWAS, the lack of explicit justification and nuance in the reporting reinforces the idea of discrete human populations.
6. PMCID 6185762. The paper earns a high rating for its exemplary handling of population descriptors in a genetic study. It correctly avoids using race as a proxy for genetic variation (Recommendation 1) and circumvents typological thinking (Recommendation 2) by employing a modern, statistically robust approach. Instead of stratifying the analysis by discrete racial or ancestral groups, the authors analyze their 'large, multi-ethnic OFC cohort' as a whole and use continuous variables to account for genetic diversity. Specifically, the paper states, 'Eighteen principal components of ancestry were identified and used as covariates in the analysis.' This method treats genetic ancestry as a continuum rather than a set of distinct categories, which is the current best practice.

The paper is also transparent about its methodology, providing a link to a full quality control report, thus fulfilling the recommendation to disclose the process of assigning labels (Recommendation 8). While the introduction references incidence rates in ‘African ancestry’, ‘European’, and ‘Asian’ populations to provide context from existing literature, the actual analysis appropriately moves away from these broad labels to a more precise, quantitative model of genetic similarity.

7. PMCID 6723657. The paper follows the recommendations extremely well. It clearly specifies the population descriptor used for inclusion, stating that participants ”self-reported as being of Taiwanese Han Chinese ancestry.” This is a transparent and appropriate use of a self-identified descriptor (Recommendations 6, 7, 8). Crucially, the paper does not use this label as a proxy for genetic variation. Instead, it correctly employs a measure of genetic similarity—principal components—to account for population structure within the cohort, stating it performed ”Adjustments for population stratification with top 10 principal components” (Recommendation 1). This method avoids typological thinking by treating genetic variation as continuous rather than categorical (Recommendation 2). The terminology is specific and avoids problematic labels (Recommendation 3)
8. PMCID 7255939. The paper earns a rating of 5 because it handles population descriptors in a manner that aligns perfectly with the NASEM recommendations. The authors state clearly and consistently that the analyses were restricted to participants of ”European origin” or ”European ancestry.” This descriptor is used for a valid methodological reason in a GWAS meta-analysis—to control for confounding due to population stratification. The authors avoid problematic terminology like ”Caucasian” (Recommendation 3). Crucially, they explicitly acknowledge the limitations of this approach and avoid typological thinking (Recommendation 2) by stating, ”Since all of the GWAS summary statistics analyzed in this study were generated from cohorts of European ancestry... the results may not be generalizable to non-European populations.” This demonstrates a clear understanding of the implications of their cohort selection and follows the recommendation to explain why descriptors were used (Recommendation 6). The paper is transparent about the source of the data and its labels, applying the descriptor consistently across the study samples (Recommendations 7 & 8).
9. PMCID 8131358. The paper largely adheres to the NASEM recommendations for the reporting of its study population. The authors are transparent and consistent in their use of the group label ‘White British’ (Recommendation 7), and they clearly state the rationale for this choice: ”due to data availability restricted the analyses to White British individuals” (Recommendations 6 & 8). Crucially, they do not use this label as a proxy for genetic variation; instead, they properly use ”the first 20 genetic principal components to control for population stratification,” which is the recommended practice for accounting for genetic similarity in a GWAS (aligning with Recommendation 1). The paper also avoids problematic terminology like ‘Caucasian’ (Recommendation 3). The rating is not a perfect 5 because, while the reporting is sound, the study design itself relies on a single, broad ancestry group, which perpetuates the lack of diversity in genomics that the NASEM report seeks to address. However, given the available data, the authors’ handling and description of their cohort are methodologically sound and transparent.

10. PMCID 8620763. The text adheres well to the recommendations by using a technical measure of genetic similarity to control for confounding rather than relying on race or ethnicity as a proxy. Specifically, it notes that the analysis adjusted for ”the first 5 ancestry principal components in INTERVAL”. This use of principal components aligns with the report’s preference for using measures of genetic similarity over discrete, typological categories (Recommendations 1 & 2). The paper also avoids problematic terminology (Recommendation 3). It does not receive a perfect score because it could provide a more comprehensive description of the ancestral makeup of the large consortia used (e.g., GECCO) to better contextualize the generalizability of its findings. However, its handling of population structure within the genetic analysis is appropriate and follows the spirit of the guidelines.

#### Supplementary Figures

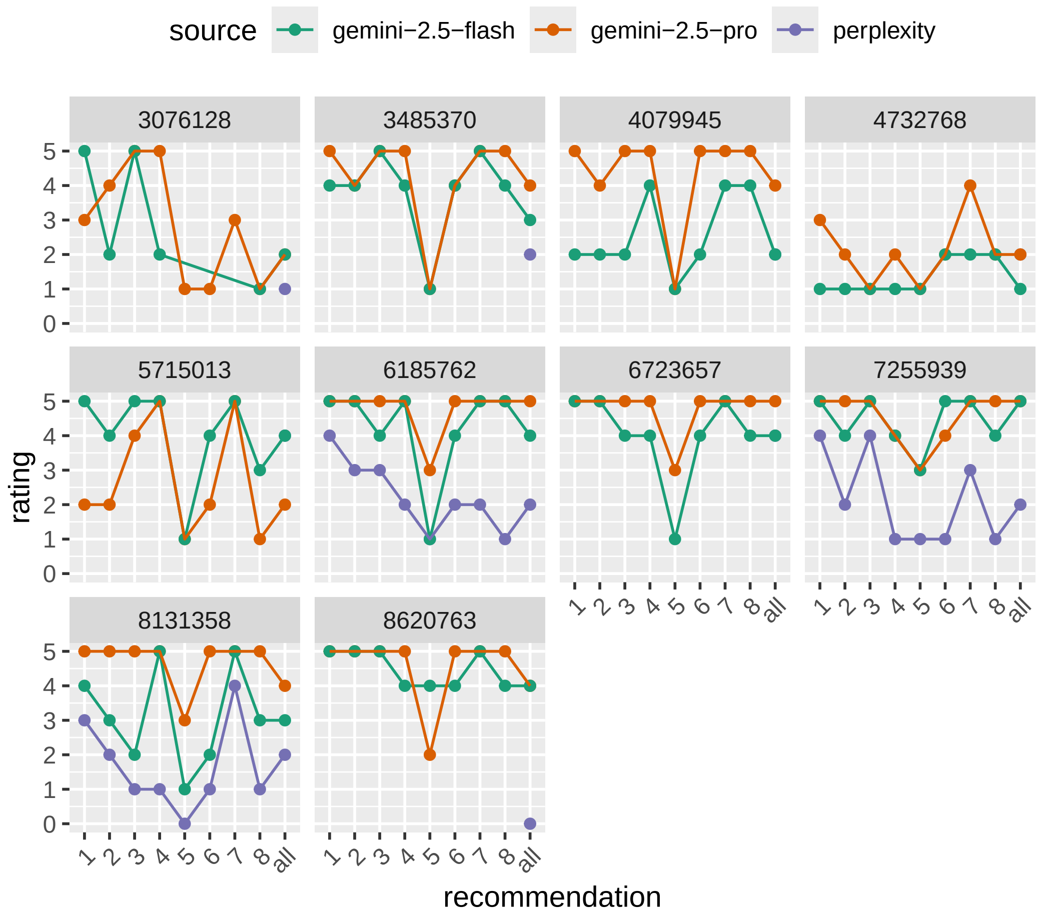


Supplementary Figure 1: Comparison of ratings from different language models for 10 papers. The PubMed Central ID of each paper is shown in the facet label. The x-axis shows the different recommendations, and the y-axis shows the rating for each recommendation.

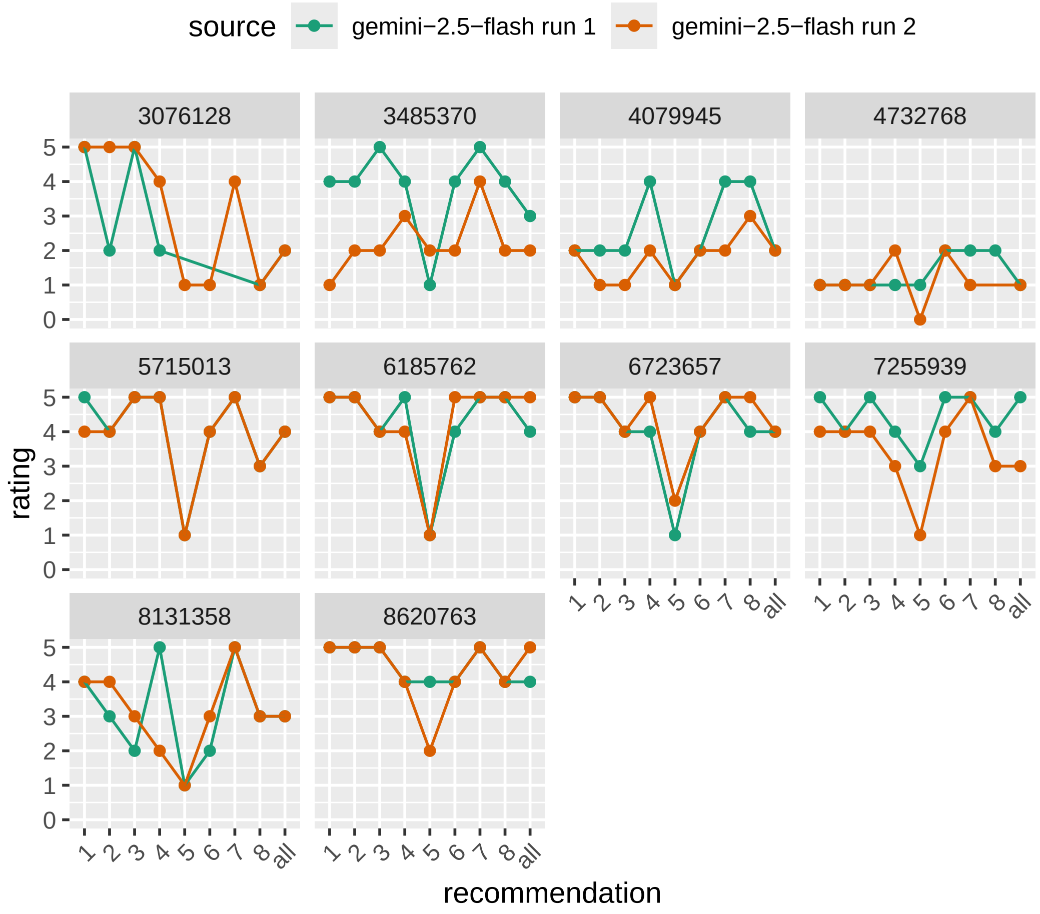


Supplementary Figure 2: Reproducability of gemini-2.5-flash ratings for 10 papers. The PubMed Central ID of each paper is shown in the facet label. The x-axis shows the different recommendations, and the y-axis shows the rating for each recommendation.

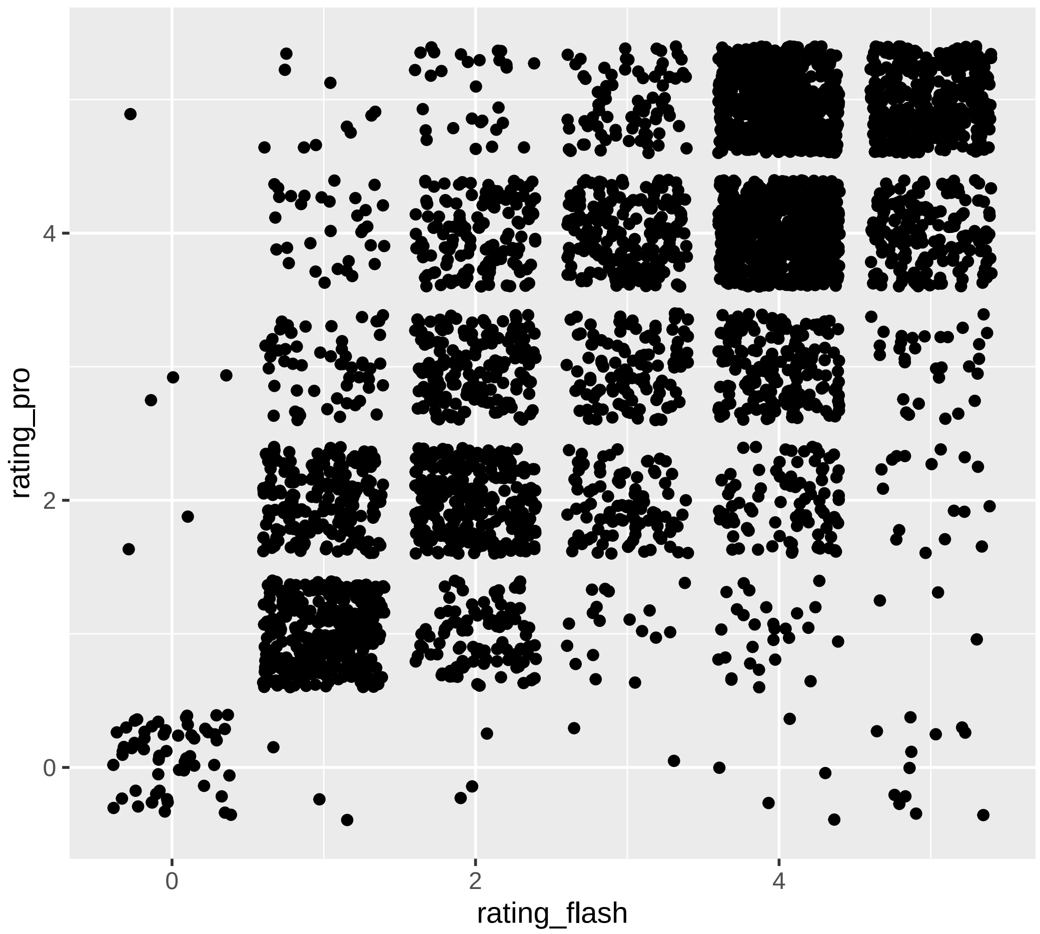


Supplementary Figure 3: Consistency of overall ratings for gemini-2.5-flash vs gemini-2.5-pro ratings for all 4007 papers. The points are jittered to show density.

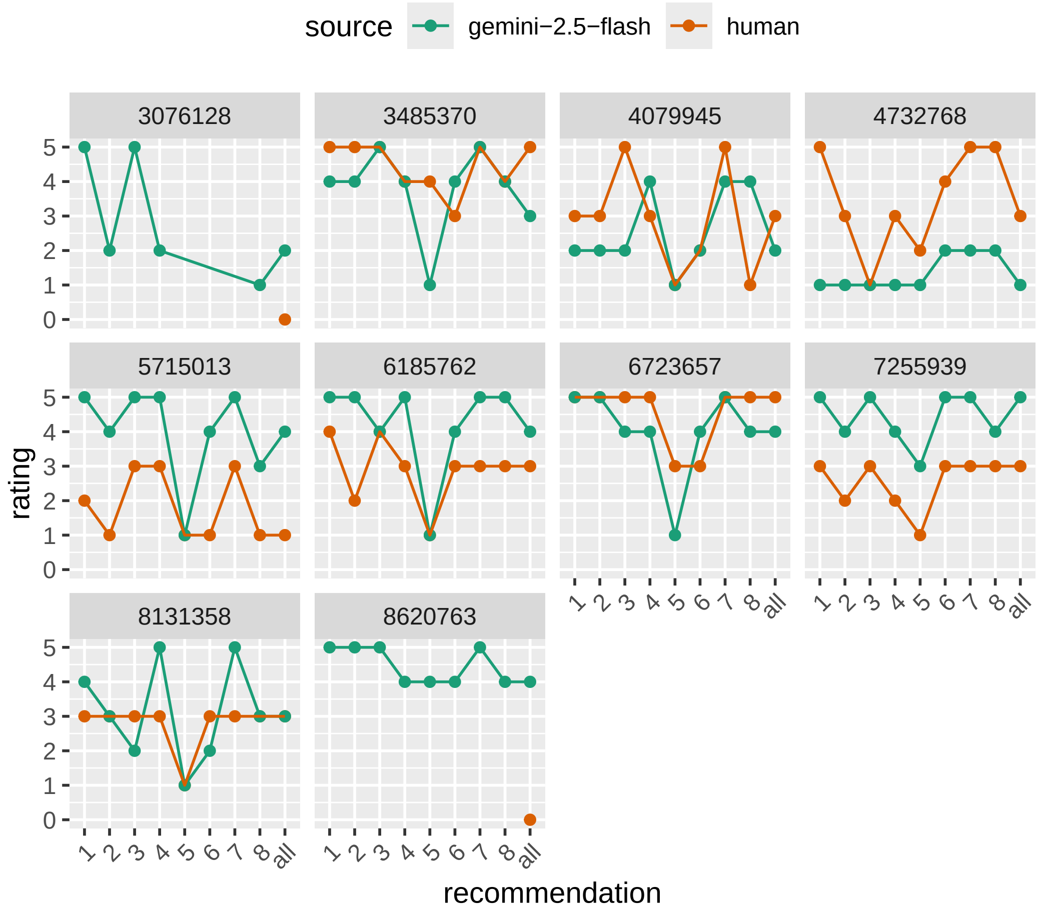


Supplementary Figure 4: Comparison of human ratings and gemini-2.5-flash ratings for 10 papers. The PubMed Central ID of each paper is shown in the facet label. The x-axis shows the different recommendations, and the y-axis shows the rating for each recommendation.

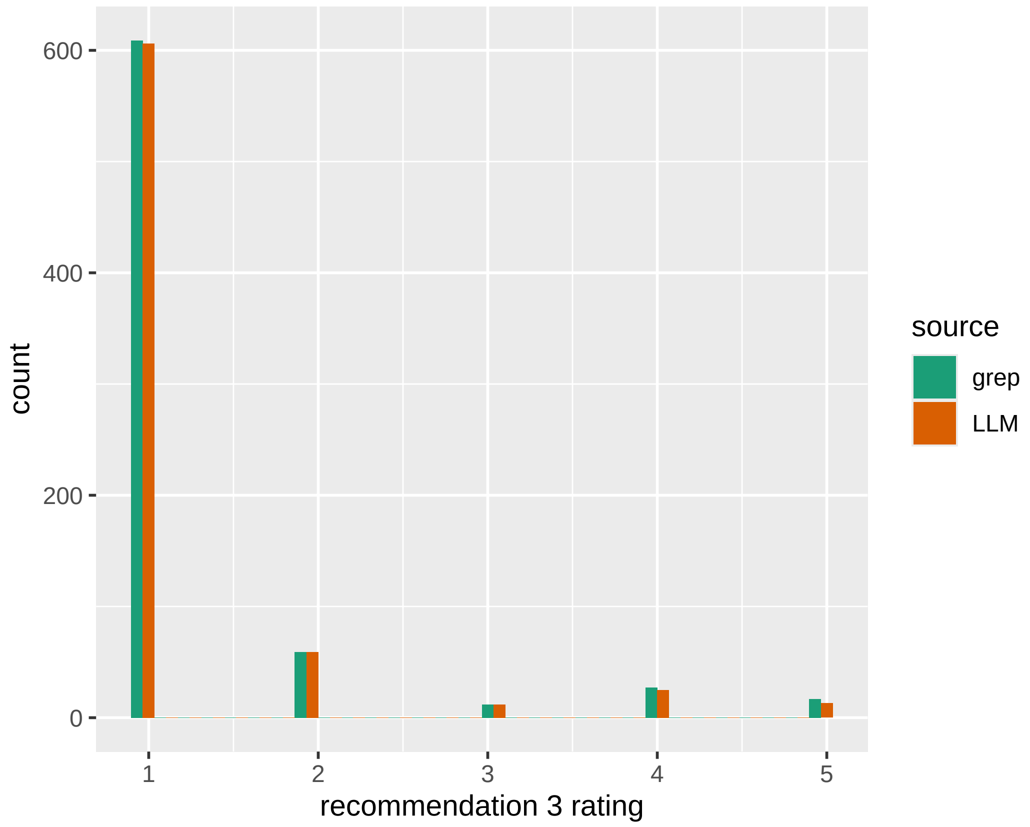


Supplementary Figure 5: Recommendation 3 ratings of papers using the term "Caucasian". Counts using both grep and the LLM are shown.

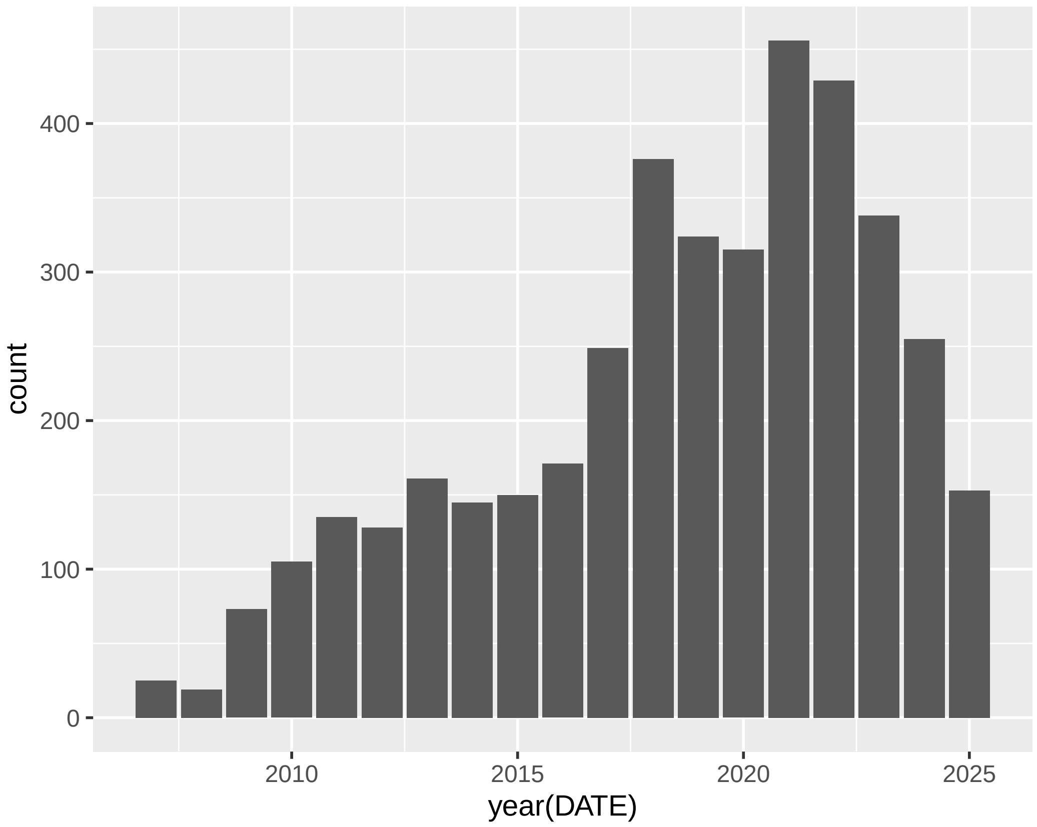


Supplementary Figure 6: Number of papers in the GWAS catalog with full text available in XML format, by year.

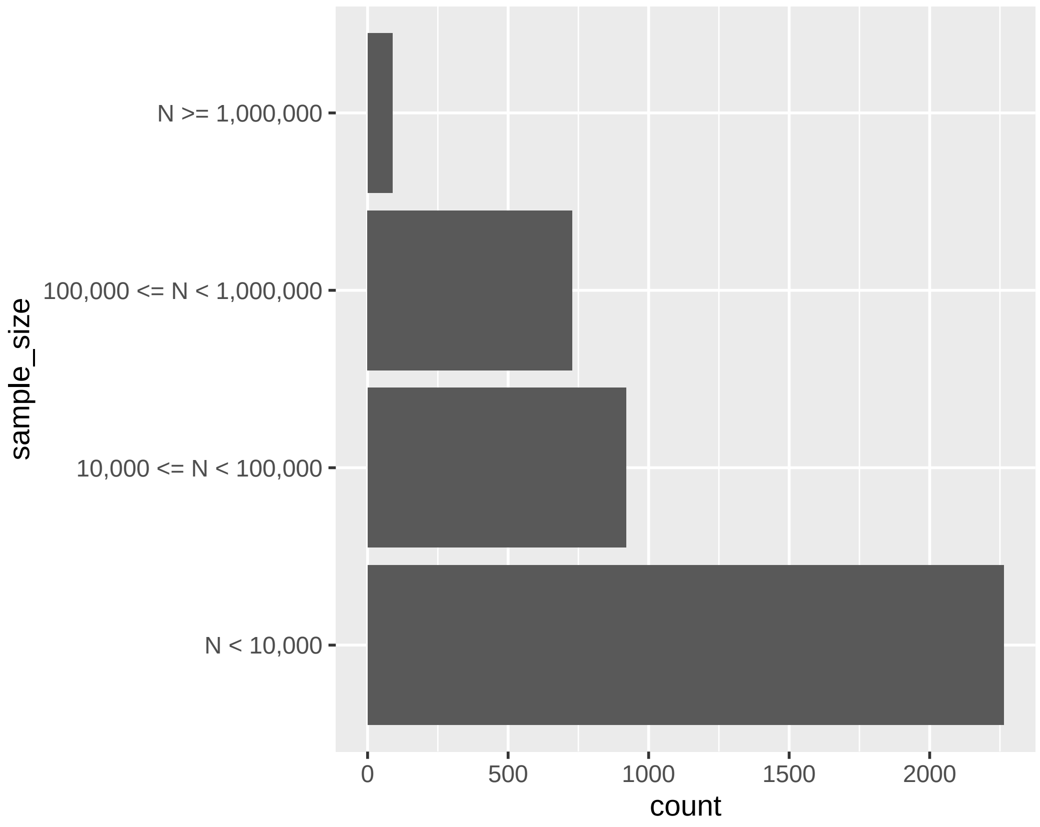


Supplementary Figure 7: Number of papers by GWAS sample size.

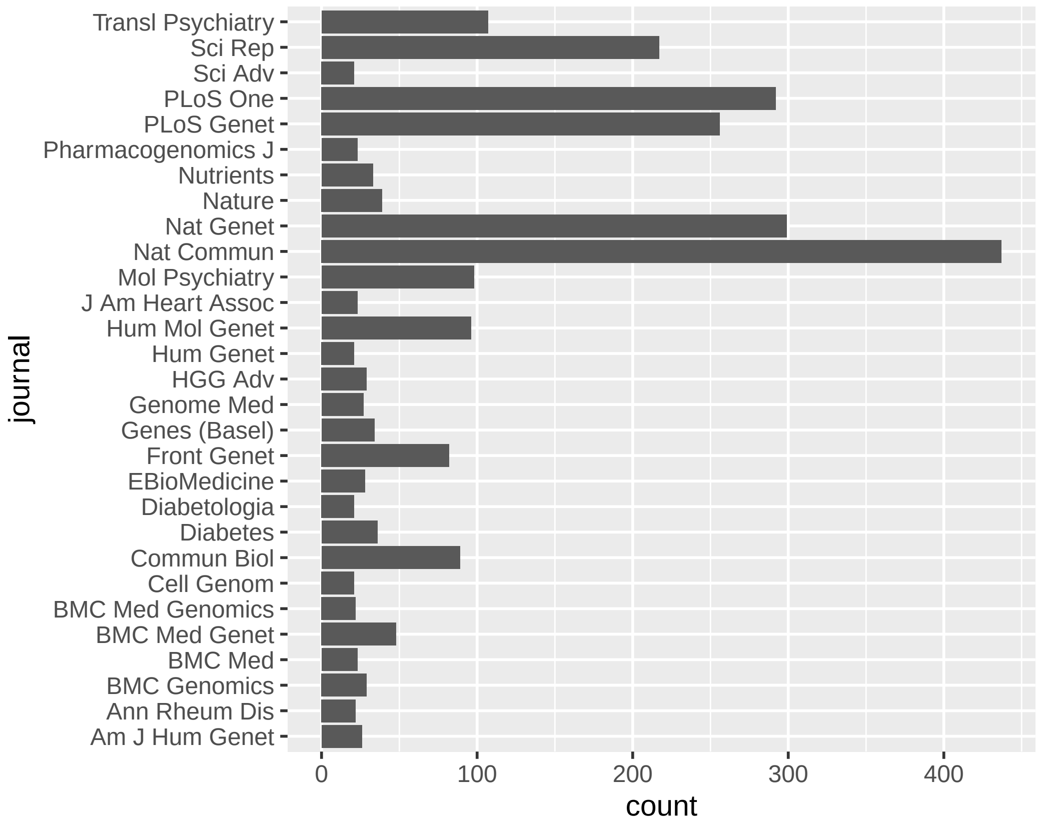


Supplementary Figure 8: Number of papers published by journal. Only journals with more than 20 papers are shown.

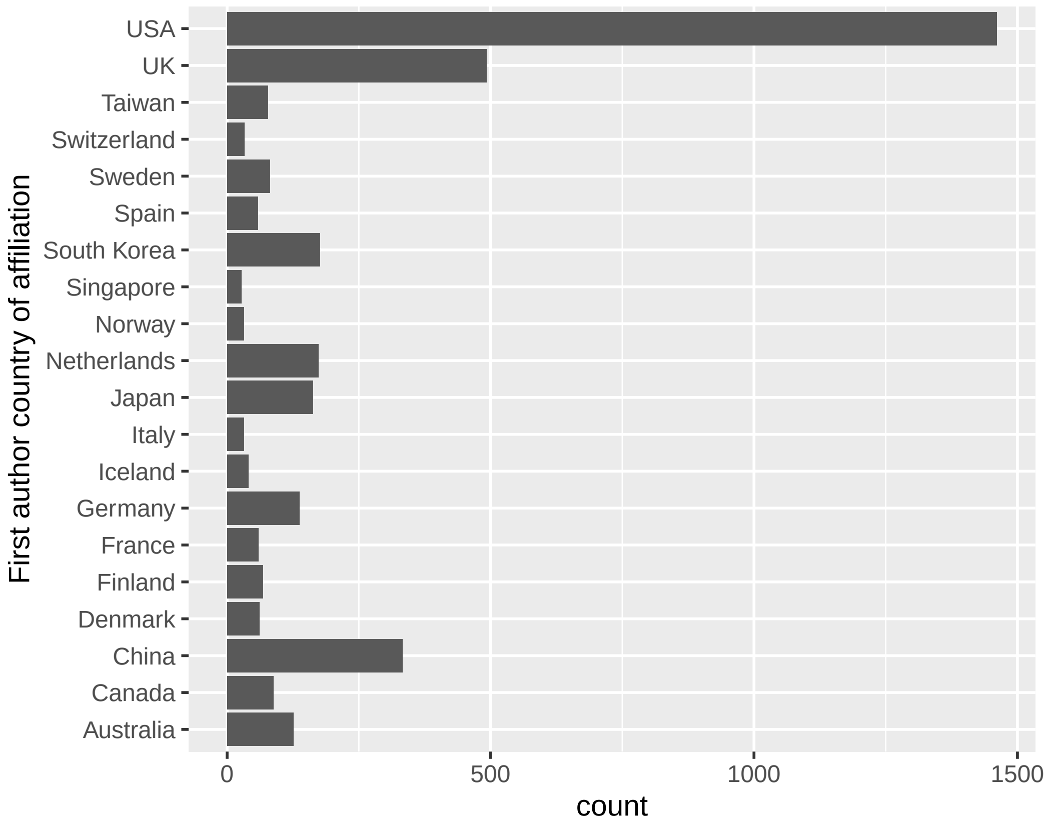


Supplementary Figure 9: Number of papers by first author's country of affiliation. Only countries with more than 20 papers are shown.

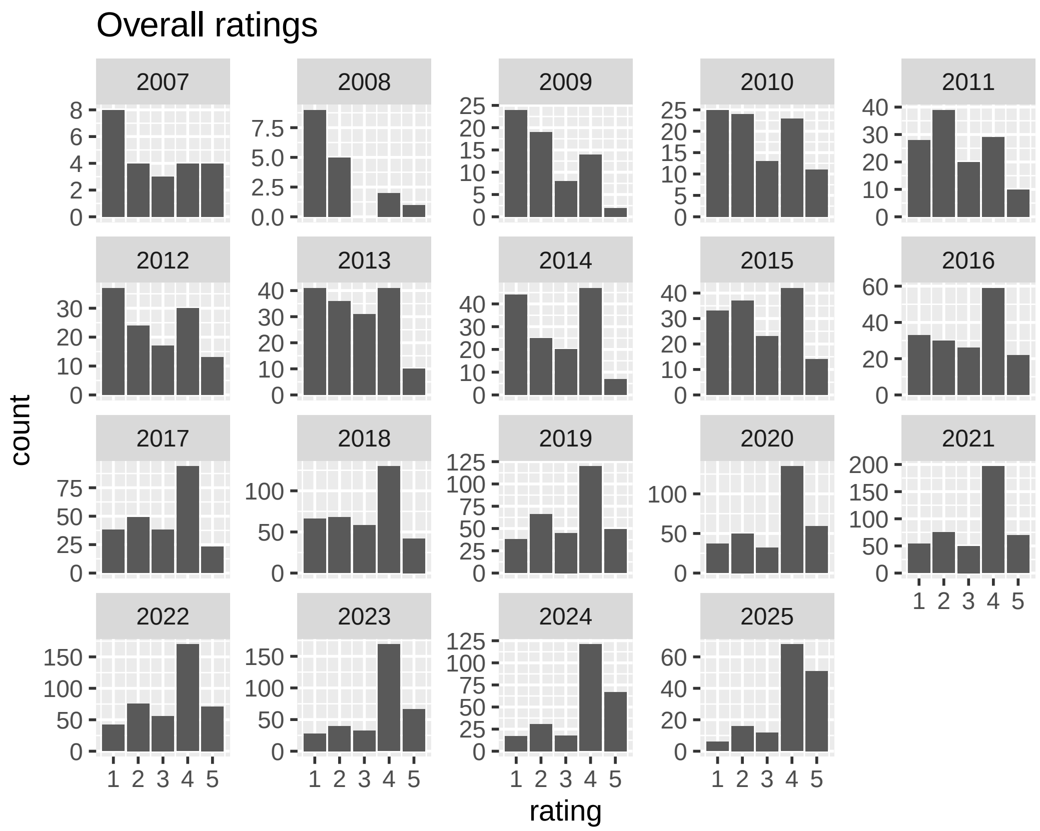


Supplementary Figure 10: Overall ratings of papers by year.

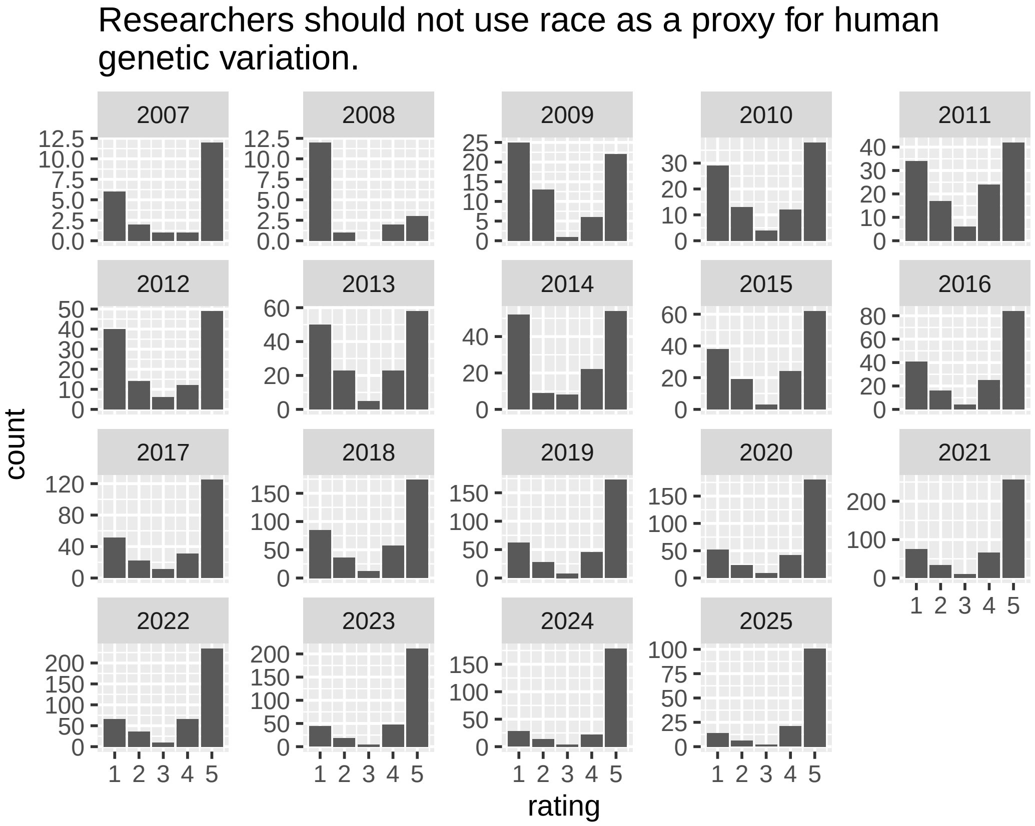


Supplementary Figure 11: Ratings for recommendation 1 by year.

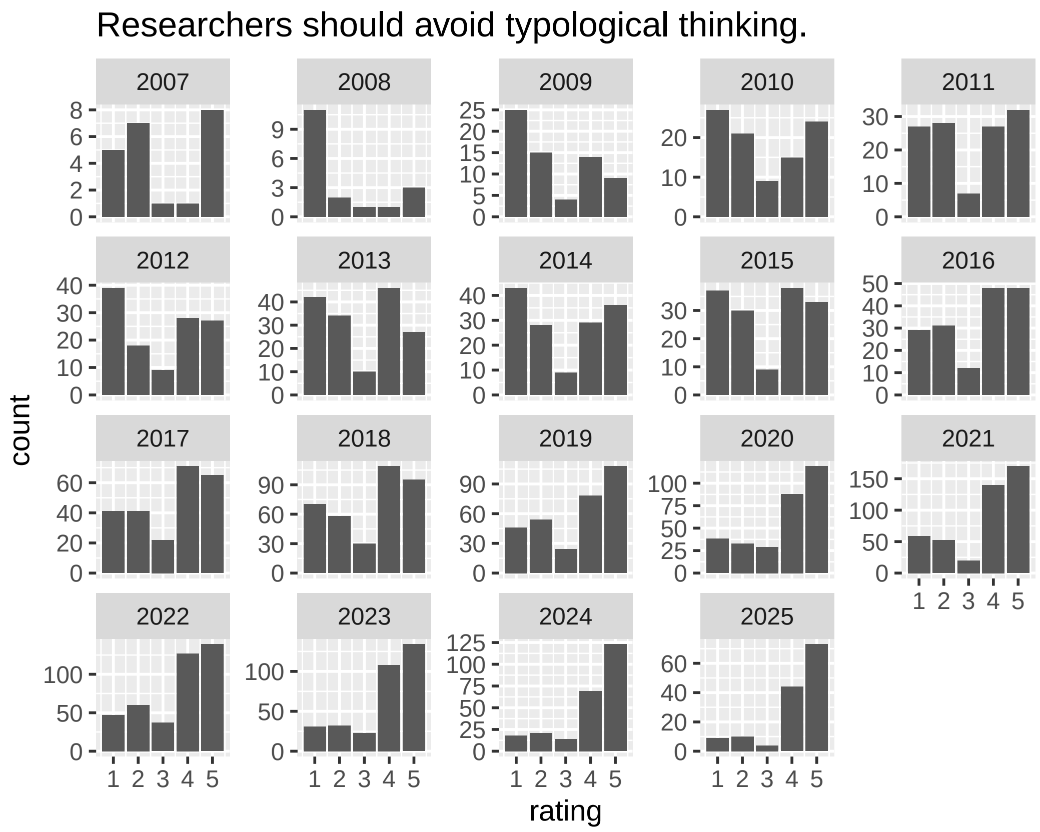


Supplementary Figure 12: Ratings for recommendation 2 by year.

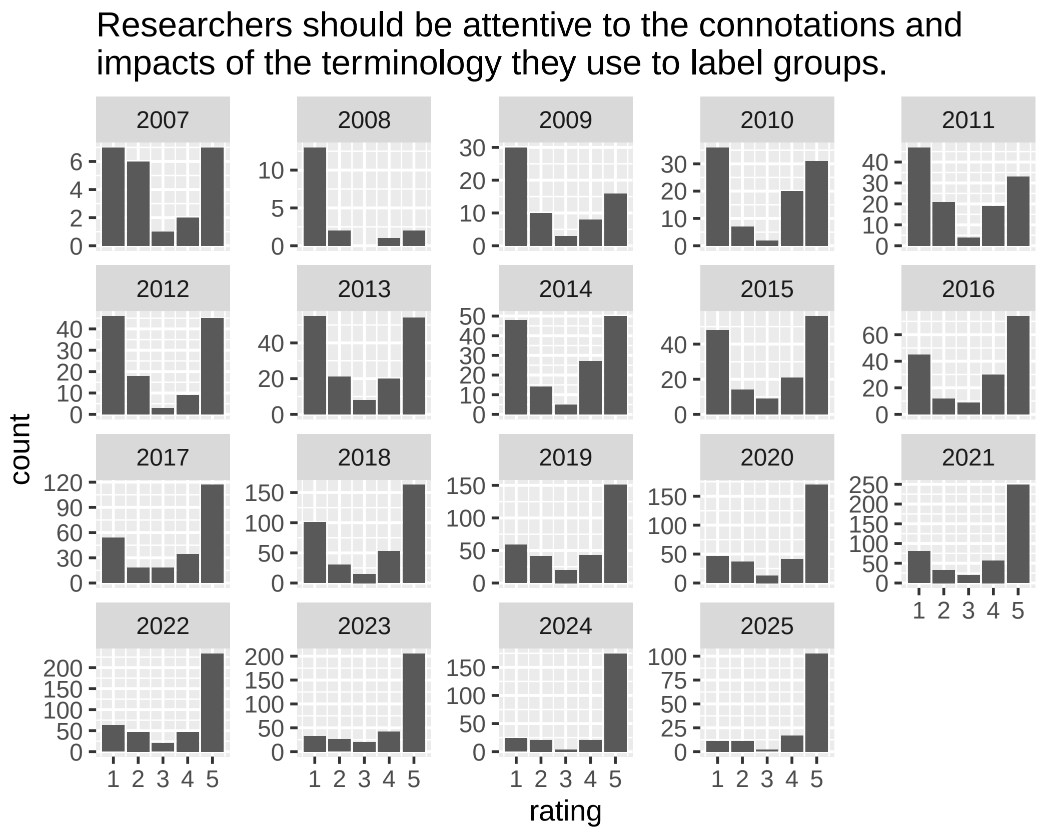


Supplementary Figure 13: Ratings for recommendation 3 by year.

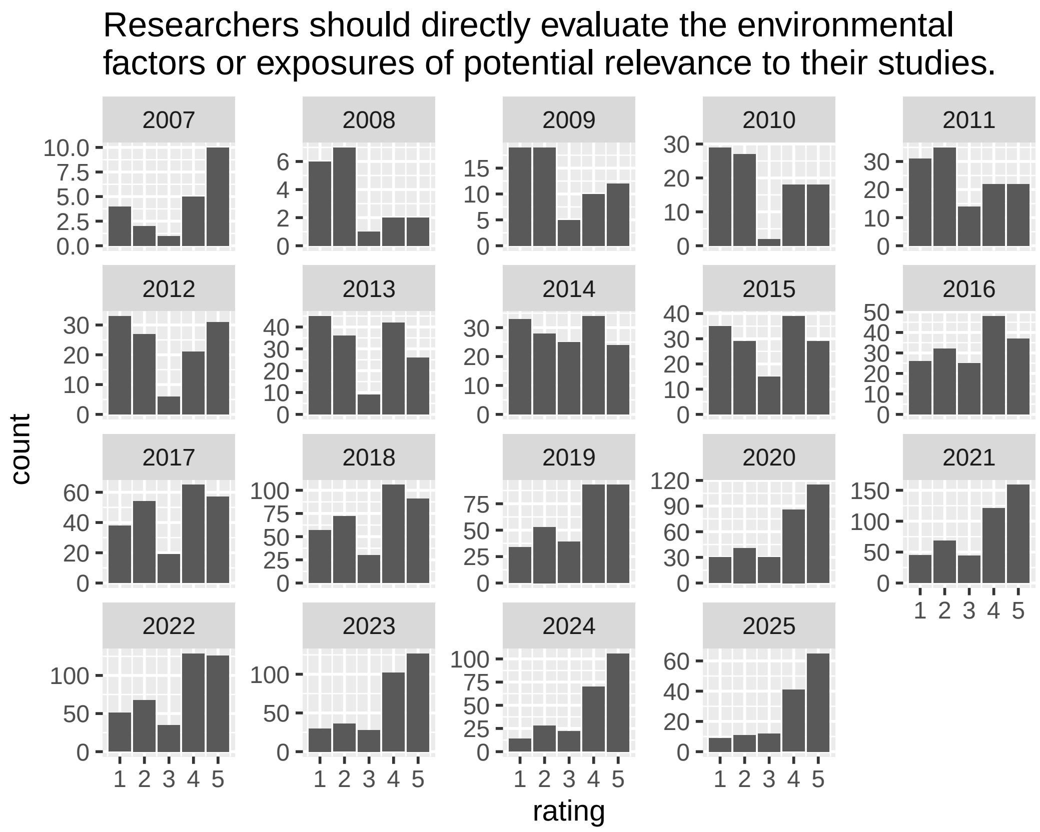


Supplementary Figure 14: Ratings for recommendation 4 by year.

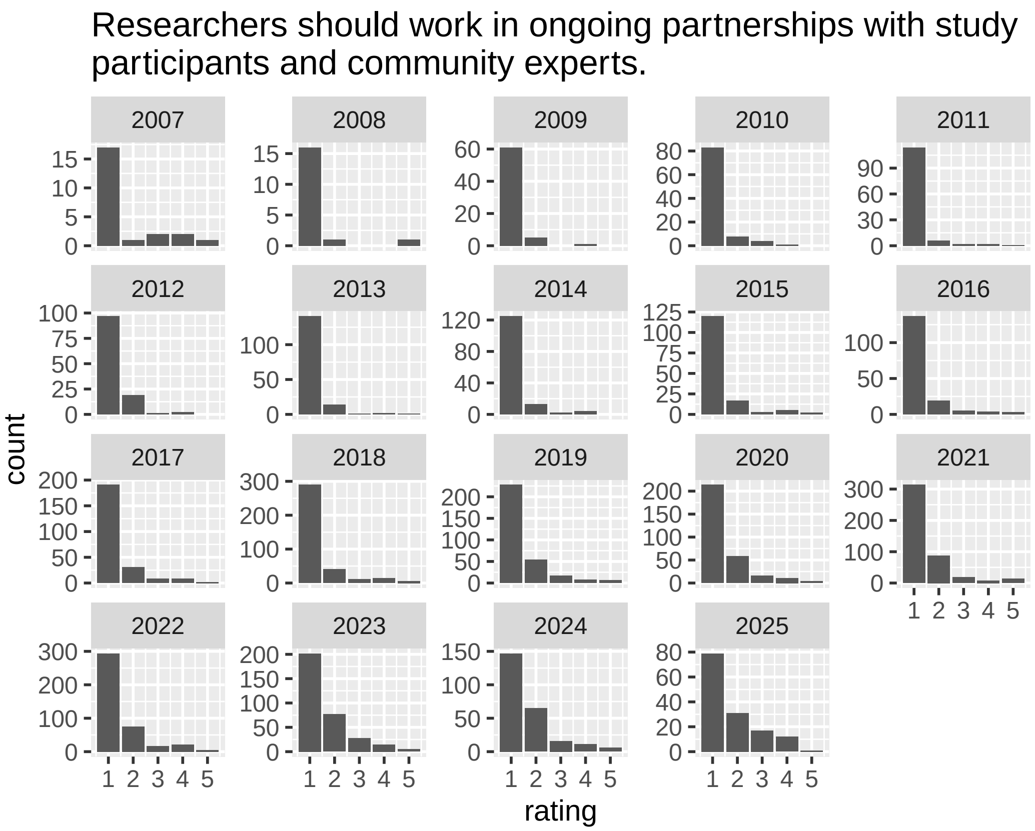


Supplementary Figure 15: Ratings for recommendation 5 by year.

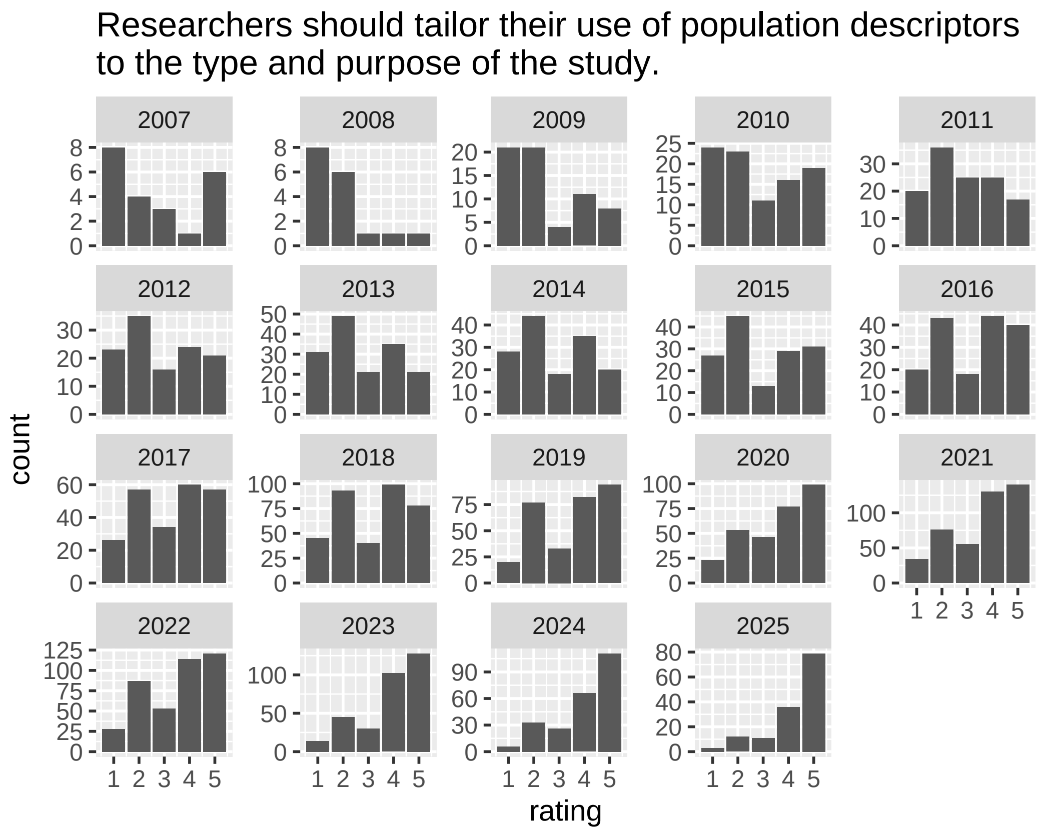


Supplementary Figure 16: Ratings for recommendation 6 by year.

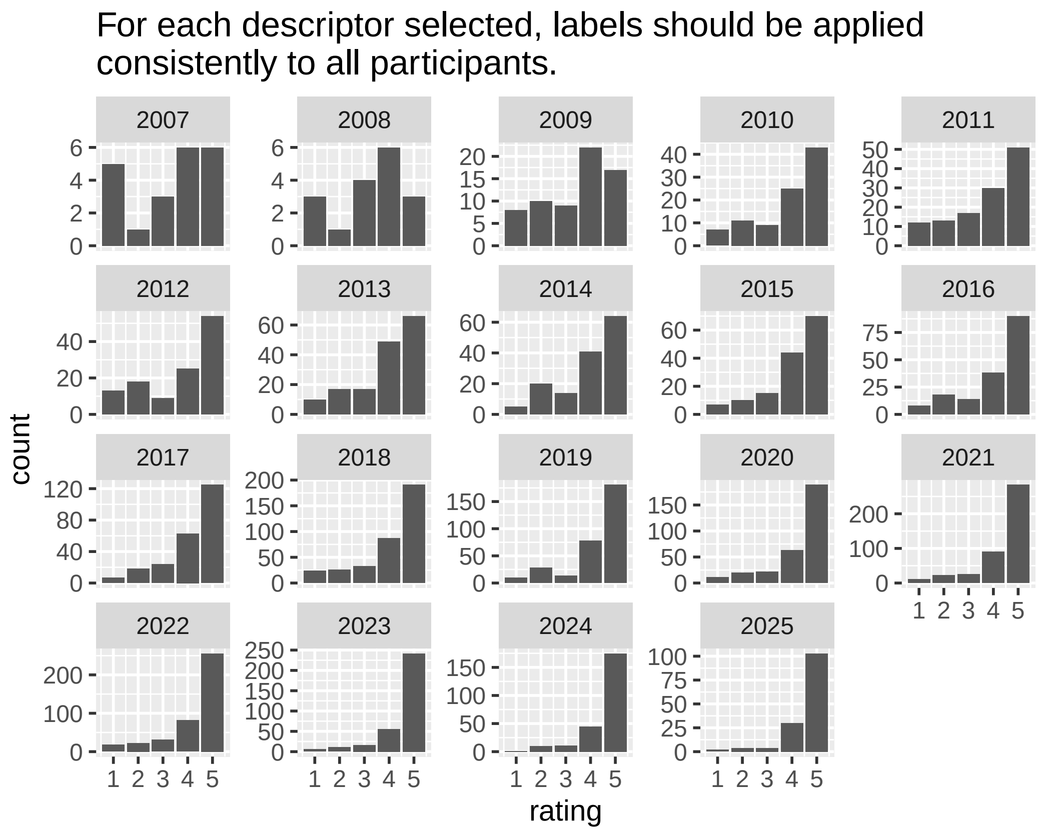


Supplementary Figure 17: Ratings for recommendation 7 by year.

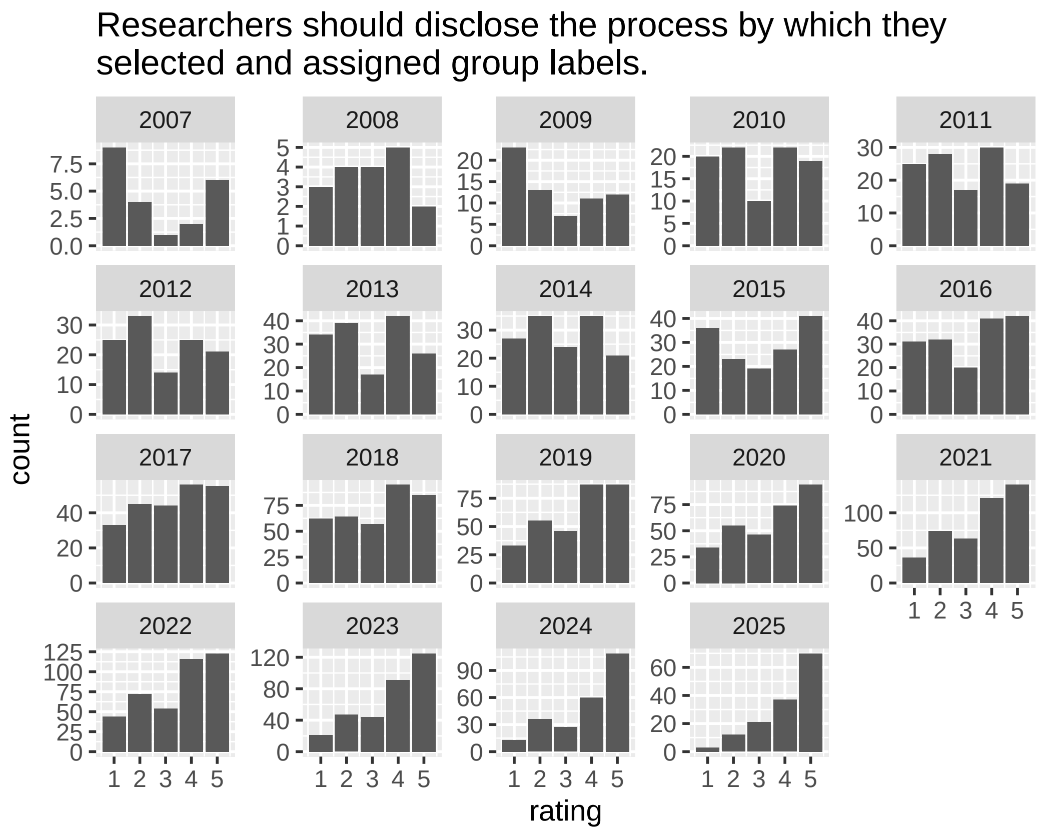


Supplementary Figure 18: Ratings for recommendation 8 by year.

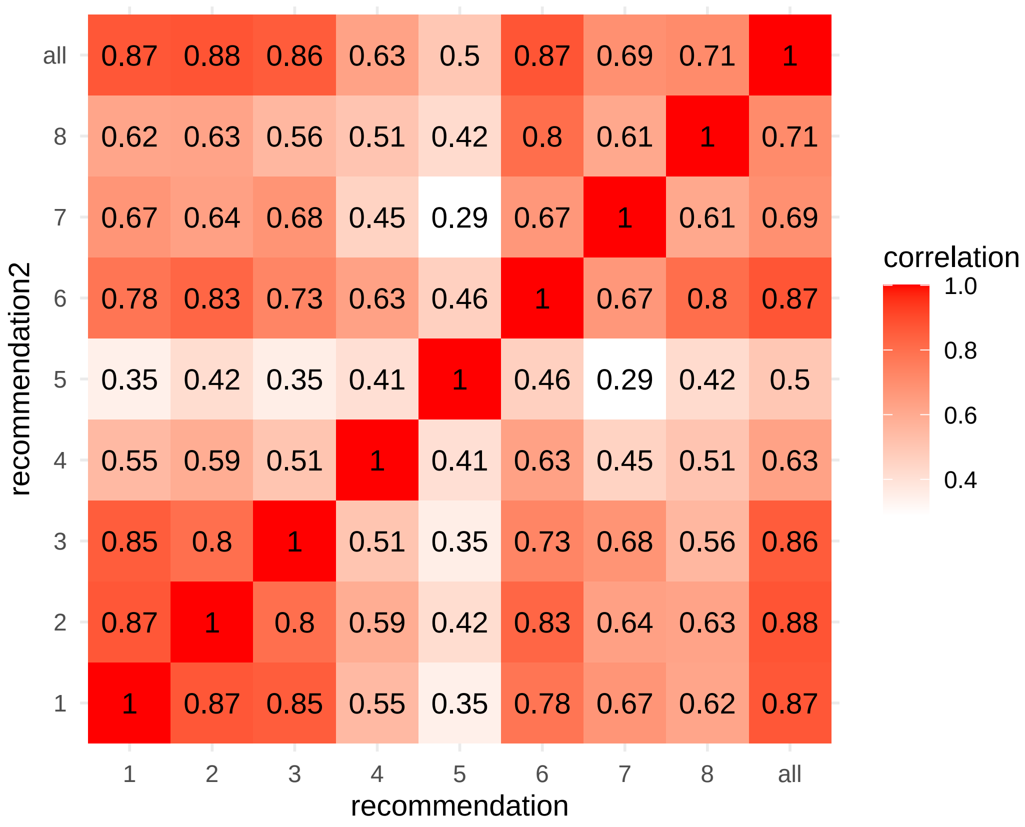


Supplementary Figure 19: Correlation structure of ratings across recommendations. "1" = "race is not genetic ancestry", "2" = "avoid typological thinking", "3" = "impacts of terminology", "4" = "use environmental factors", "5" = "work with communities", "6" = "tailor to study purpose", "7" = "apply labels consistently", "8" = "disclose labeling process", "all" = "overall".

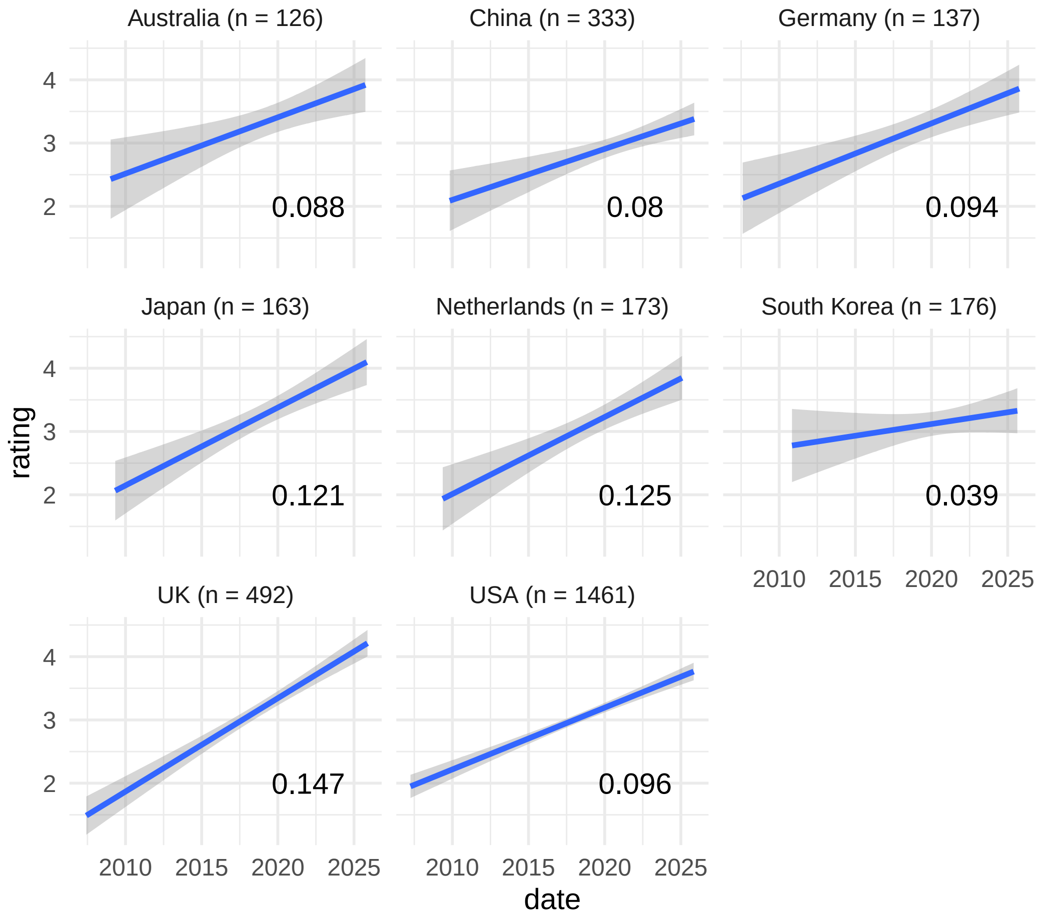


Supplementary Figure 20
